# Single-cell profiling uncovers a *Muc4*-expressing metaplastic gastric cell type sustained by *Helicobacter pylori*-specific inflammation

**DOI:** 10.1101/2022.12.20.521287

**Authors:** Valerie P. O’Brien, Yuqi Kang, Meera K. Shenoy, Greg Finak, William C. Young, Julien Dubrulle, Lisa Koch, Armando E. Rodriguez Martinez, Jazmine A. Snow, Jeffery Williams, Elizabeth Donato, Surinder K. Batra, Cecilia C.S. Yeung, Susan Bullman, Meghan A. Koch, Raphael Gottardo, Nina R. Salama

## Abstract

Mechanisms for *Helicobacter pylori* (*Hp*)-driven stomach cancer are not fully understood. In a transgenic mouse model of gastric preneoplasia, concomitant *Hp* infection and induction of constitutively active KRAS (*Hp*+KRAS+) alters metaplasia phenotypes and elicits greater inflammation than either perturbation alone. Gastric single-cell RNA-seq showed that *Hp*+KRAS+ mice had a large population of metaplastic pit cells that expressed the intestinal mucin *Muc4* and the growth factor amphiregulin. Metaplastic pit cells were associated with macrophage and T cell inflammation and prevented by gastric immunosuppression. Lineage tracing showed that *Muc4* was not dependent on cell-intrinsic KRAS activity, and lineage-derived cells had a limited propensity for growth as organoids, demonstrating that metaplastic pit cells are largely not self-renewing. Finally, MUC4 expression was significantly associated with proliferation in human gastric cancer samples. These studies identify an *Hp*-associated metaplastic pit cell lineage, also found in human gastric cancer tissues, whose expansion is driven by *Hp*-dependent inflammation.

**Statement of Significance:** Using a mouse model, we have delineated metaplastic pit cells as a pre-cancerous cell type whose expansion requires *H. pylori*-driven inflammation. In humans, metaplastic pit cells show enhanced proliferation as well as enrichment in precancer and early cancer tissues, highlighting an early step in the gastric metaplasia to cancer cascade.

## Introduction

About 20-25% of cancer deaths are due to chronic inflammation-associated carcinomas, which follow a developmental cascade wherein tissue injury causes persistent inflammation that leads to metaplasia (conversion of one normal cell type to another), dysplasia (presence of abnormal cells) and finally cancer ^1^. Targeting inflammation is an attractive strategy to prevent or treat inflammation-associated cancers, but the nature of the immunopathology is variable and must be elucidated for each cancer type.

Gastric adenocarcinoma is a canonical example of chronic inflammation-associated carcinoma. About 80% of cases are attributable to infection with *Helicobacter pylori* (*Hp*) ^2^, a bacterium that colonizes the stomach of half the world’s population ^3^. *Hp* infection causes gastric inflammation (superficial gastritis). In some individuals, inflammation triggers the loss of gastric acid-producing parietal cells (chronic atrophic gastritis) and subsequent elevation of stomach pH. In response to parietal cell loss, digestive enzyme-producing chief cells (hypothesized to be a cell of origin of gastric metaplasia ^4, 5^) can transdifferentiate into metaplastic cells, which can precede the appearance of dysplasia and gastric cancer ^6–12^. Thus, *Hp* infection elicits the chronic inflammation that promotes gastric adenocarcinoma development, and eradication of *Hp* with antibiotics significantly reduces risk of gastric cancer development and recurrence ^13–15^.

However, the exact mechanism(s) through which *Hp* infection and associated inflammation drive gastric disease remain poorly understood.

In mice, key aspects of human preneoplastic progression, including the development of metaplasia and mild dysplasia, can be modeled within 12 weeks by induction of a constitutively active *Kras* allele in gastric chief cells ^4^. We previously found that concomitant *Hp* infection of these KRAS transgenic mice (*Hp*+KRAS+) elicited greater dysplasia compared to *Hp*-KRAS+ mice ^16^. This phenotype could be due to either *Hp* infection <u>accelerating</u> the disease trajectory that results from constitutively active KRAS, or *Hp* infection driving a <u>different</u> disease trajectory. To distinguish between these possibilities, we used scRNA-seq to examine changes in cell populations that occur in the context of gastric metaplasia with and without *Hp* infection. We found that mice with concomitant chronic *Hp* infection and transgene-driven gastric metaplasia developed an abundant population of *Muc4*-expressing metaplastic pit cells that exhibited a genetic signature of inflammation and epithelial-to-mesenchymal transition, compared to mice without *Hp* infection. *Hp*-specific gastric inflammation was required: gastric immunosuppression and antibiotic eradication of *Hp* were each independently protective, whereas gastric infection with another bacterium, *Fusobacterium nucleatum*, did not elicit severe inflammation or pit cell metaplasia. Studies of human samples revealed that pit cells acquire the expression of intestinal mucins as disease develops, and MUC4 expression is significantly associated with cell proliferation. Taken together, these studies define a novel cell state that is driven by *Hp*- mediated inflammation and associated with cell proliferation and gastric cancer.

## Results

### scRNA-seq revealed heterogeneous pit cell populations associated with *Hp* infection and metaplasia induction

We previously found that in *Mist1-CreERT2, LSL-Kras*^G12D^ (“*Mist1-Kras”*) mice, concomitant *Hp* infection and metaplasia driven by constitutively active KRAS induction led to altered gastric gland architecture, changes to metaplasia marker expression, increased dysplasia and severe inflammation within 12 weeks ^16^. To obtain an unbiased view of changes to gastric cell type abundance and gene expression, we conducted single cell RNA-sequencing (scRNA-seq) with mice +/- *Hp*, +/- KRAS at 12 weeks (**Figure 1A, Table S1**). We also analyzed tissues from KRAS+ mice +/- *Hp* at 6 weeks to explore how cell populations changed over time. Because constitutively active KRAS is targeted to the corpus (body) of the stomach in this model, the gastric forestomach and antrum were discarded prior to sample processing. This experiment yielded 22,050 cells, >75% of which were viable based on mitochondrial gene filtering. Cluster analyses followed by visualization with UMAP (Uniform Manifold Approximation and Projection) identified 25 cell clusters, which we manually annotated based on marker gene expression (UMAP #1, **Figure 1B, Figure S1-2, Table S2**). The gene expression profiles from the two samples collected at six weeks were largely similar to those from samples collected at 12 weeks (**Figure S1-2**). We detected several cell lineages, including putative immune cells (e.g., macrophages and T cells) as well as non-hematopoietic cells (e.g., endothelial and muscle cells). However, we first investigated gastric epithelial cells to address our overarching question of whether *Hp*+KRAS+ mice had an altered or accelerated disease trajectory compared to *Hp*- KRAS+ mice. Glands in the gastric corpus are comprised of a variety of epithelial cell types, including acid-producing parietal cells, digestive enzyme-producing chief cells, and two mucus-producing cell types: pit (foveolar) cells at the surface and neck cells in the middle of the glands. Each of these cell types were found in a large epithelial “megacluster” (**Figure 1B**, dashed lines). We observed three clusters within this megacluster that expressed the mucin and classical pit cell marker *Muc5Ac,* which we annotated as pit_1, pit_2 and pit_3.

**Figure 1.**
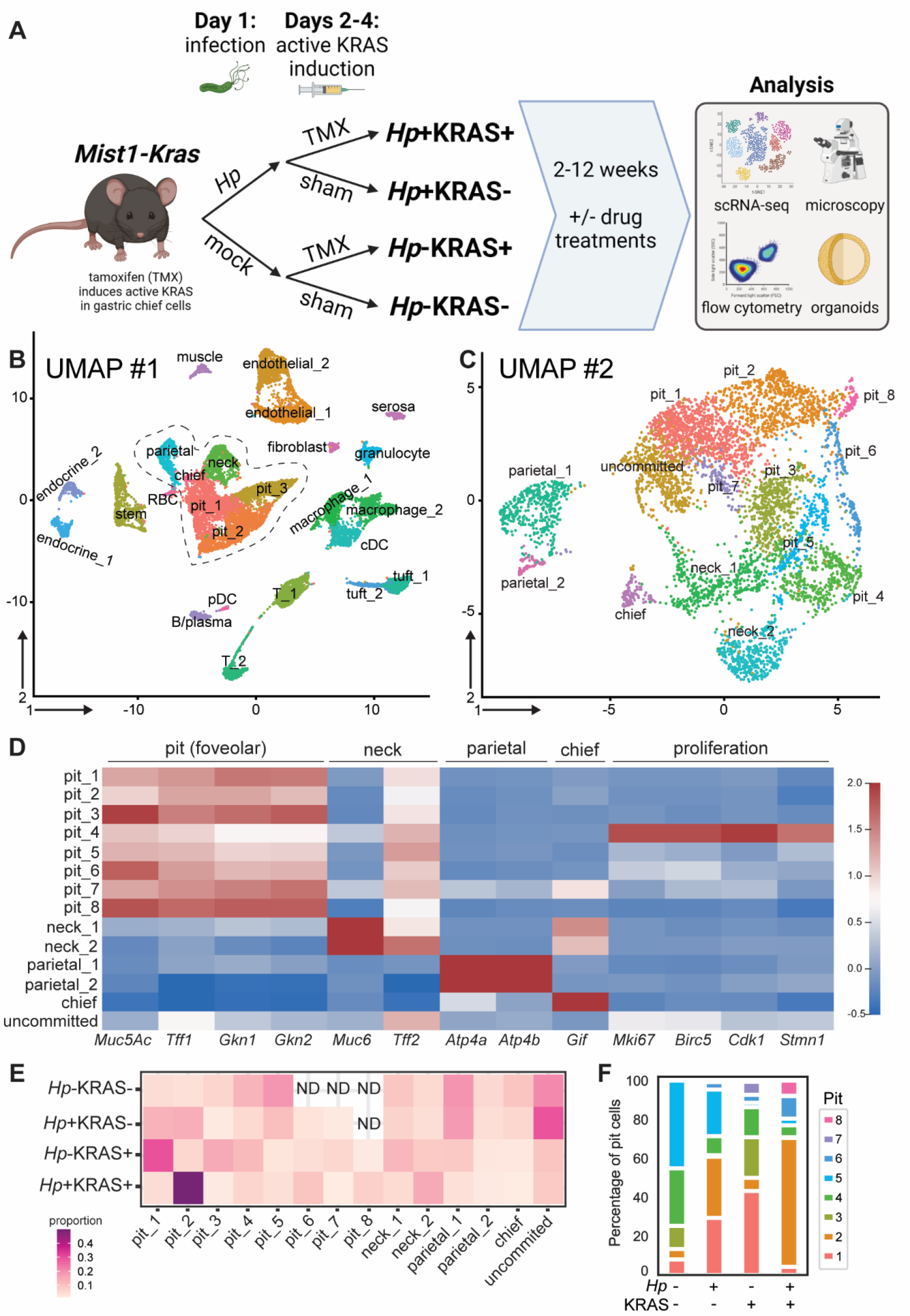
Gastric scRNA-seq reveals expansion of pit cell populations in *Hp*+KRAS+ mice. A) Overview and experimental timeline for mouse experiments, created with BioRender.com. TMX, tamoxifen. **B-F)** Mice +/- *Hp* infection and +/- constitutively active KRAS induction were used for gastric scRNA-seq. Corpus single cell suspensions were prepared from: one *Hp*-KRAS+ and one *Hp*+KRAS+ mouse at six weeks; and at 12 weeks, three *Hp*-KRAS- and two *Hp*+KRAS- (each pooled into one sample), one individual (never cryopreserved) and two pooled *Hp*-KRAS+ mice, and one individual (never cryopreserved) and two pooled *Hp*+KRAS+ mice. **B)** Cluster analyses followed by UMAP (Uniform Manifold Approximation and Projection) visualization of the entire dataset yielded 25 clusters, including epithelial and immune cell types, which were manually annotated based on gene expression. Endocrine, enteroendocrine cell; cDC, conventional dendritic cell; RBC, erythrocyte/reticulocyte; pDC, plasmacytoid dendritic cell. **C)** Cells from the 12 week time point in the central epithelial cell megacluster, outlined with the dotted line in **B**, were subjected to reclustering followed by UMAP visualization, yielding 14 clusters that were manually annotated based on gene expression. **D)** The normalized expression of marker genes for the indicated gastric epithelial cell types is shown for the 14 epithelial clusters from UMAP #2 in **C**. **E)** The proportion of each of the 14 epithelial clusters in the indicated treatment groups at 12 weeks is shown (rows add to 100%). ND, cell cluster was not detected in the indicated treatment group. **F)** The distribution of pit cell clusters within each treatment group in **E** is shown.

To better discriminate among these gastric epithelial cell types and compare their abundance across all treatment groups, we re-clustered the cells in the central epithelial megacluster from the 12 week time point alone. This analysis yielded 14 clusters that we manually annotated based on gene expression (UMAP #2, **Figure 1C, Table S3**). Most of the clusters expressed low to moderate levels of the trefoil factor *Tff2*, which is expressed by several gastric corpus epithelial cell types ^17^ (**Figure 1D**). One cluster expressed *Tff2* but no other known marker genes, suggesting it may be an uncommitted or progenitor cell type. Further analysis showed that the expression of specific ribosomal genes differentiated this cluster (**Figure S3, Table S4**). Eight of the clusters expressed classical pit cell genes (*Muc5Ac*, trefoil factor *Tff1* and the gastrokines *Gkn1* and *Gkn2*) and we annotated them as pit_1 through pit_8 (**Figure 1D**). The pit_4 cluster also expressed proliferation markers (*Mki67*, *Birc5*, *Cdk1* and *Stmn1*), suggesting these cells may be pit cell progenitors.

The abundance of the different epithelial clusters varied in the different treatment groups (**Figure 1E**, **S4**). Parietal and chief cells were less abundant in *Hp*-KRAS+ mice and least abundant in *Hp*+KRAS+ mice, consistent with our previous analysis of tissues from these mice^16^. To explore pit cell heterogeneity, we determined the proportion of all pit cells annotated as each individual cluster (**Figure 1F**). The predominant pit cell clusters in each treatment group were: pit_5 (45%) and pit_4 (30%) in *Hp*-KRAS- (healthy) mice; pit_2 (32%) and pit_1 (29%) in *Hp*+KRAS- mice; pit_1 (43%) in *Hp*-KRAS+ mice; and pit_2 (66%) in *Hp*+KRAS+ mice. Thus, pit cells are a heterogeneous population whose composition dynamically changes due to infection and/or metaplasia.

### *Hp*+KRAS+ mice have an abundant population of *Muc4*-expressing pit cells

Because pit cell populations differed between mice with and without metaplasia, we queried whether any pit cell clusters expressed metaplasia-associated genes (**Figure 2A**). The neck_2 cluster expressed several genes associated with a type of gastric adenocarcinoma-associated metaplasia called spasmolytic polypeptide-expressing metaplasia (SPEM) ^18–20^, including gastrokine *Gkn3*, the chloride channel *Cftr*, clusterin (*Clu*), *Wfcd2* (HE4), the cell surface glycoprotein *Cd44* and the water channel protein *Aqp5*, but had low expression of the SPEM markers *Mal2* (a proteolipid family member) and the non-coding RNA *Xist*. This cluster was most abundant in *Hp*+KRAS+ mice (**Figure 1E, S4**), which we previously showed have increased SPEM compared to *Hp*-KRAS+ mice at 12 weeks ^16^. We next investigated the expression of intestinal metaplasia (IM) genes. Along with the neck_2 cluster, pit_2, pit_6 and pit_8 stood out from the other clusters due to low or moderate expression of several IM-related genes: the polymeric immunoglobulin receptor *Pigr*, keratin *Krt20* and villin *Vil1*. The IM and candidate tumor suppressor glycoprotein *Dmbt1* was expressed by these clusters and several additional pit cell clusters, and the IM marker trefoil factor *Tff3* was expressed in pit_8 cells. The intestinal metaplasia-associated mucin *Muc2* was not expressed in any cell type, but another intestinal mucin, *Muc4*, was highly expressed in the pit_8 cluster and moderately in pit_2, pit_6 and neck_2. Finally, *Tacstd2* (TROP2), a dysplasia and stem cell marker ^21^, was not detected in any of the UMAP #2 cells, though it was detected in the stem cell cluster in UMAP #1 (**Figure S5**).

**Figure 2.**
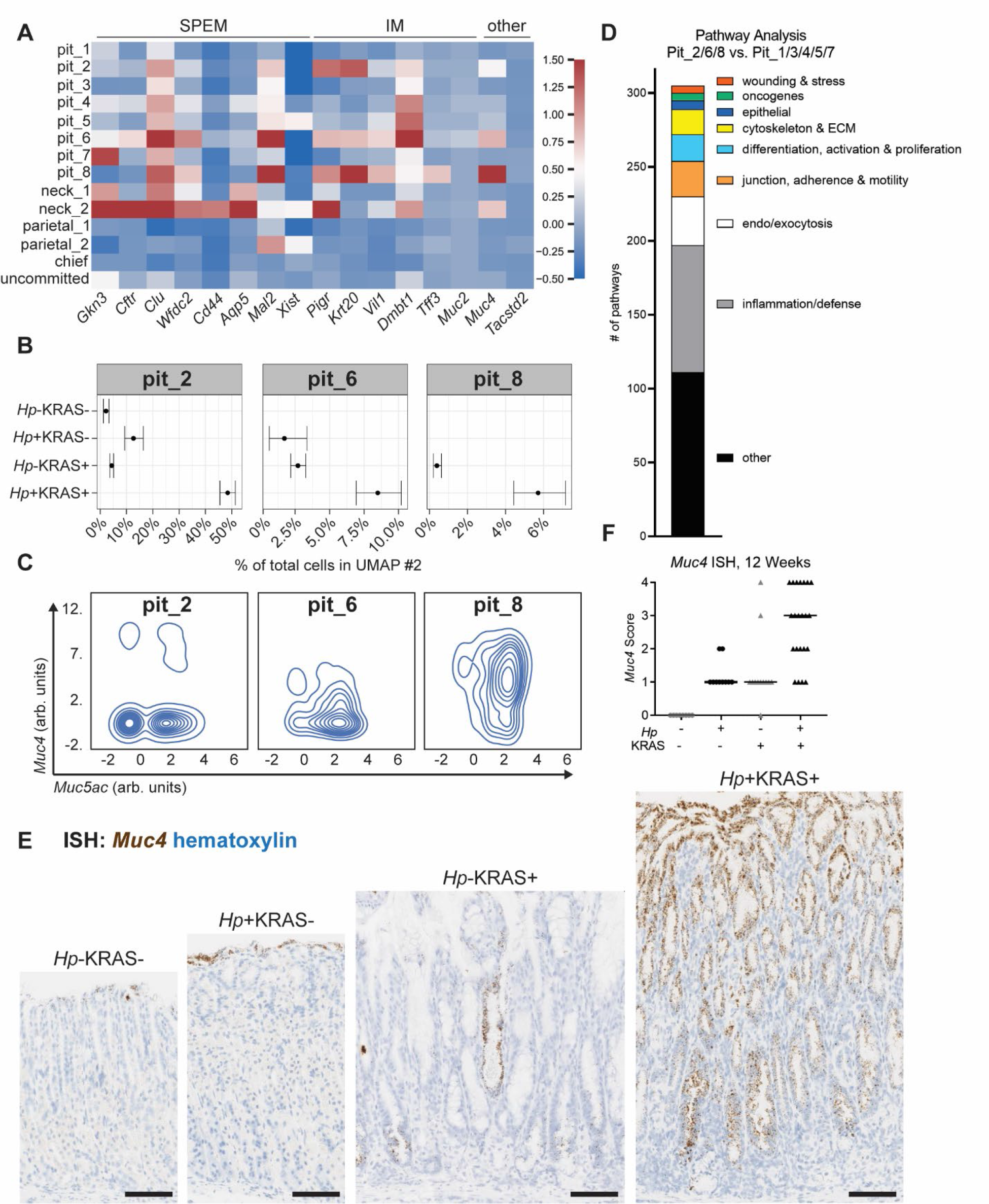
*Hp*+KRAS+ mice have abundant metaplastic pit cells that express the intestinal mucin *Muc4*. A) The normalized expression of the indicated metaplasia and dysplasia genes is given for the cells from UMAP #2 (Figure 1C). **B)** The proportion of cells assigned to the indicated pit cell clusters is shown. Error bars represent the confidence interval that a given percentage of cells would be identified as the given cluster type based on their observed distribution. **C)** The contour plots show *Muc5Ac* (x-axis) and *Muc4* (y-axis) expression in the indicated pit cell clusters from UMAP #2. Expression is given as arbitrary units and negative values are due to data scaling for visualization purposes. **D)** Gene set enrichment analysis was performed to assess biological pathways and processes enriched in pit_2, pit_6 and pit_8 cells relative to the five other pit cell clusters, using Hallmark, KEGG and GO pathway datasets from the MSigDB database. **E-F)** *In situ* hybridization (ISH) was used to detect the intestinal mucin *Muc4* (brown) in gastric corpus tissue (counterstained with hematoxylin, blue) from the indicated treatment groups at 12 weeks. **E)** Representative images are shown. Scale bars, 100 µm. **F)** Tissues were scored in a blinded fashion. N=5 independent experiments were performed with n=3-8 mice per group. Data represent actual values from each individual mouse and bars indicate the median values.

The pit_2, pit_6 and pit_8 clusters were all expanded in *Hp*+KRAS+ mice (**Figure 1F, 2B**); pit_6 and pit_8 were not detected in healthy mice. We detected *Muc4* expression in 8% of cells in pit_2, 36% in pit_6 and 84% in pit_8 (**Figure 2C**). To probe what biological processes and pathways differentiate these clusters from the other pit-related clusters, we performed gene set enrichment analysis with Gene Ontology (GO), KEGG and Hallmark databases, comparing pit_2, pit_6 and pit_8 to the other five pit cell clusters (**Figure 2D, Table S5**). A total of 305 pathways and processes were enriched in pit_2/6/8 cells, which we categorized based on function (**Figure 2D**). The most common categorized pathways (n=85) pertained to inflammation and host defense, including interferon response, TNFα signaling via NFκB, complement, response to cytokines and IL-2/STAT5 signaling. As well, 33 pathways pertained to endocytosis/exocytosis. Another 24 pertained to cell-cell junctions, adherence and motility and 17 pertained to the cytoskeleton and extracellular matrix, suggesting that pit_2/6/8 cells may be undergoing cellular remodeling. Eighteen pathways pertained to cell differentiation, activation or proliferation. Other enriched pathways of interest pertained to epithelial cells (six pathways including epithelial-mesenchymal transition [EMT], a hallmark of cancer development) and wound healing and stress (five pathways), which could suggest that these cells are generated by, or in response to, epithelial damage. Finally, five pathways involving gastric cancer-associated oncogenes were upregulated: KRAS signaling, the P53 pathway and ErbB signaling. Because pit_2, pit_6 and pit_8 cells were most abundant in *Hp*+KRAS+ mice and expressed metaplasia-associated genes and pathways related to EMT and oncogenesis, we termed these populations metaplastic pit cells.

To explore metaplastic pit cell localization and expansion, we focused on *Muc4*, which encodes an intestinal goblet cell mucin that normally is expressed in cells of the first gland adjacent to the forestomach-glandular stomach junction and sparsely at the luminal surface of the glandular stomach in mice ^22, 23^. *In situ* hybridization (ISH) showed robust *Muc4* expression throughout the corpus of *Hp*+KRAS+ mice at 12 weeks (**Figure 2E**), which we confirmed using spatial transcriptomics (**Figure S6**). *Muc4*-expressing pit cells were also found in *Hp*+KRAS+ mice at six weeks by scRNA-seq and ISH (**Figure S7A-B**). We thus used *Muc4* expression as a read- out for metaplastic pit cells. In a blinded semi-quantitative scoring of mice at 12 weeks (see Methods), *Muc4* expression was greater in *Hp*+KRAS+ mice than in any other group (**Figure 2F**). *Muc4* expression did not correlate with gastric *Hp* bacterial burden (**Figure S7C**). In summary, the expansion of *Muc4*-expressing metaplastic pit cells requires the combination of *Hp* infection and active KRAS; neither *Hp* infection nor metaplasia induction alone are sufficient to drive this phenotype within the 12 week experimental time frame.

### Metaplastic pit cell development requires sustained, *Hp*-specific gastric inflammation

Because metaplastic pit cells were enriched for transcripts associated with immune-related pathways, we explored whether inflammation was required for the increased accumulation of these cells. It was previously shown that group 2 innate lymphoid cells could promote gastric metaplasia as a wound-healing response after tissue injury ^24^. However, the type 2 cytokines *Il4*, *Il5*, *Il9* and *Il13* were not detected in our scRNA-seq dataset (not shown) and type 2 inflammatory pathways were not uncovered in our gene set enrichment analysis (**Table S5**).

Thus, we performed flow cytometry (**Figure S8-9**) to profile immune cell populations in the gastric lamina propria at six weeks, a time point at which metaplasia and proliferation phenotypes in *Hp*+KRAS+ mice have already begun to diverge from *Hp*-KRAS+ mice (**Figure S7** and ^16^). We compared *Hp*+KRAS+ mice to *Hp*+KRAS- and *Hp*-KRAS+ mice (**Figure 3A-H**), reasoning that the immune cells involved in metaplastic pit cell accumulation should only be increased in *Hp*+KRAS+ mice. Neutrophils were elevated in both *Hp*+KRAS+ and *Hp*+KRAS- mice compared to *Hp*-KRAS+ mice (**Figure 3A**), indicating that their accumulation was due to *Hp* infection alone. However, circulating monocytes, macrophages and conventional dendritic cells were all significantly increased in *Hp*+KRAS+ mice compared to the control groups (**Figure 3B-D**). Among lymphocytes, we saw increased cytotoxic (CD8+) and helper (CD4+) T cells in *Hp*+KRAS+ mice. The majority of these cells were CD103+, consistent with a tissue-resident phenotype, and had high expression of CD44, consistent with an effector phenotype (**Figure 3E-F**). As well, *Hp*+KRAS+ mice had the greatest numbers of regulatory T cells (**Figure 3G**). Thus, *Hp*+KRAS+ mice have significant increases in gastric lamina propria myeloid cells and effector T cells, but not neutrophils.

**Figure 3.**
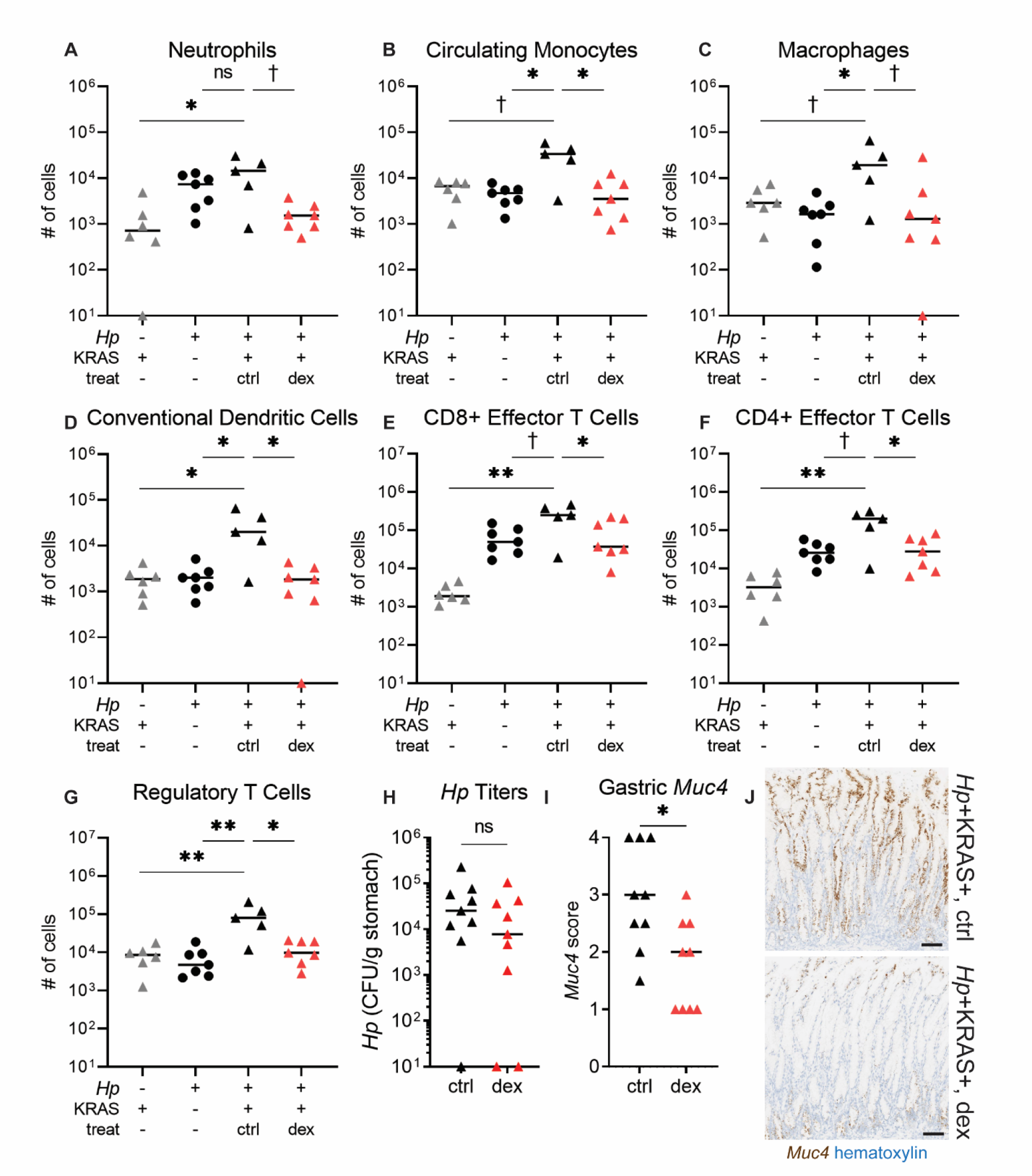
Gastric immunosuppression prevents variant pit cell expansion. A-G) Flow cytometry was performed at six weeks using *Hp*-KRAS+, *Hp*+KRAS-, and *Hp*+KRAS+ mice who received either oral dexamethasone (‘dex’) starting at two weeks or water alone (control, ‘ctrl’). Graphs indicate the frequency of live leukocytes isolated from the gastric lamina propria of the indicated mouse groups. **A)** CD4-CD8-Ly6C+Ly6G+ neutrophils, **B)** CD11b variable Ly6C+ MHC II- circulating monocytes, **C)** CD64+F4/80+ macrophages, **D)** CD11c high MHC II high conventional dendritic cells, **E)** CD3+CD8α+CD103+CD44+ effector T cells, **F)** CD3+CD4+ CD103+CD44+ effector T cells, **G)** CD3+CD4+CD103+CD44+FoxP3+ regulatory T cells. N=2 independent experiments were conducted and data from one experiment is shown. Data points represent actual values for each individual mouse, zeroes are plotted at the limit of detection (10 cells), and bars indicate median values. **H-J)** *Hp*+KRAS+ mice that were control-treated or dexamethasone-treated are shown. **H)** *Hp* was cultured from stomach homogenate supernatants at time of euthanasia. CFU, colony-forming units. Zeroes are plotted at the limit of detection (10 CFU). **I-J)** *Muc4* (brown) was detected by ISH in N=2 independent experiments with n=3-6 mice per group. **I)** *Muc4* expression was scored in a blinded fashion. **J)** Representative images are shown. Scale bars, 100 µm. † *P* < 0.10, * *P* ≤ 0.05, ** *P* < 0.01, ns, not significant, Mann-Whitney U test.

To test whether inflammation was necessary for metaplastic pit cell expansion, we treated *Hp*+KRAS+ mice with the immunosuppressive glucocorticoid dexamethasone. Mice received dexamethasone *ad libitum* in the drinking water starting at two weeks and were euthanized at six weeks. *Hp* bacterial loads were not significantly different between the two groups (**Figure 3H**). As expected, dexamethasone treatment significantly reduced the accumulation of gastric immune subsets compared to control-treated *Hp*+KRAS+ mice (**Figure 3A-G**). Importantly, gastric *Muc4* expression was also significantly reduced in dexamethasone-treated mice (**Figure 3I-J**). Thus, gastric inflammation is necessary for metaplastic pit cell development in *Hp*+KRAS+ mice and is primarily comprised of increased gastric macrophages and T cells.

We wondered whether gastric inflammation and metaplastic pit cells might develop in response to bacterial infection as a general phenomenon. To test this hypothesis, we infected mice with the Gram-negative bacterium *Fusobacterium nucleatum* (*Fn*), an oral commensal species that is also significantly associated with gastrointestinal cancers ^25, 26^. Because we were unable to reproducibly colonize the stomachs of healthy mice with *Fn* (data not shown), we introduced *Fn* into KRAS+ animals. Six weeks after induction of constitutively active KRAS, mice were gavaged with *Fn* or PBS as a control. At 12 weeks, *Fn* was recovered from the stomachs of all infected mice, though it colonized less efficiently than *Hp* did (median 2x10^2^ colony-forming units [CFU] per g tissue compared to 5x10^3^ CFU/g for *Hp* ^16^, **Figure S10**). We found that gastric Muc4 expression was not increased beyond control levels in *Fn*-infected mice, highlighting a unique relationship between *Hp*-induced inflammation and metaplasia in our mouse model (**Figure 4A**). To confirm that *Hp* was responsible for inducing inflammation that drove metaplastic pit cell expansion, we tested samples from *Hp*+KRAS+ mice given antibiotics from six to eight weeks post infection and constitutively active KRAS induction ^16^. *Muc4* expression was strongly reduced at 12 weeks in antibiotic-treated mice (**Figure 4A**), again supporting our conclusion that metaplastic pit cell expansion requires sustained, *Hp*-specific inflammation.

**Figure 4.**
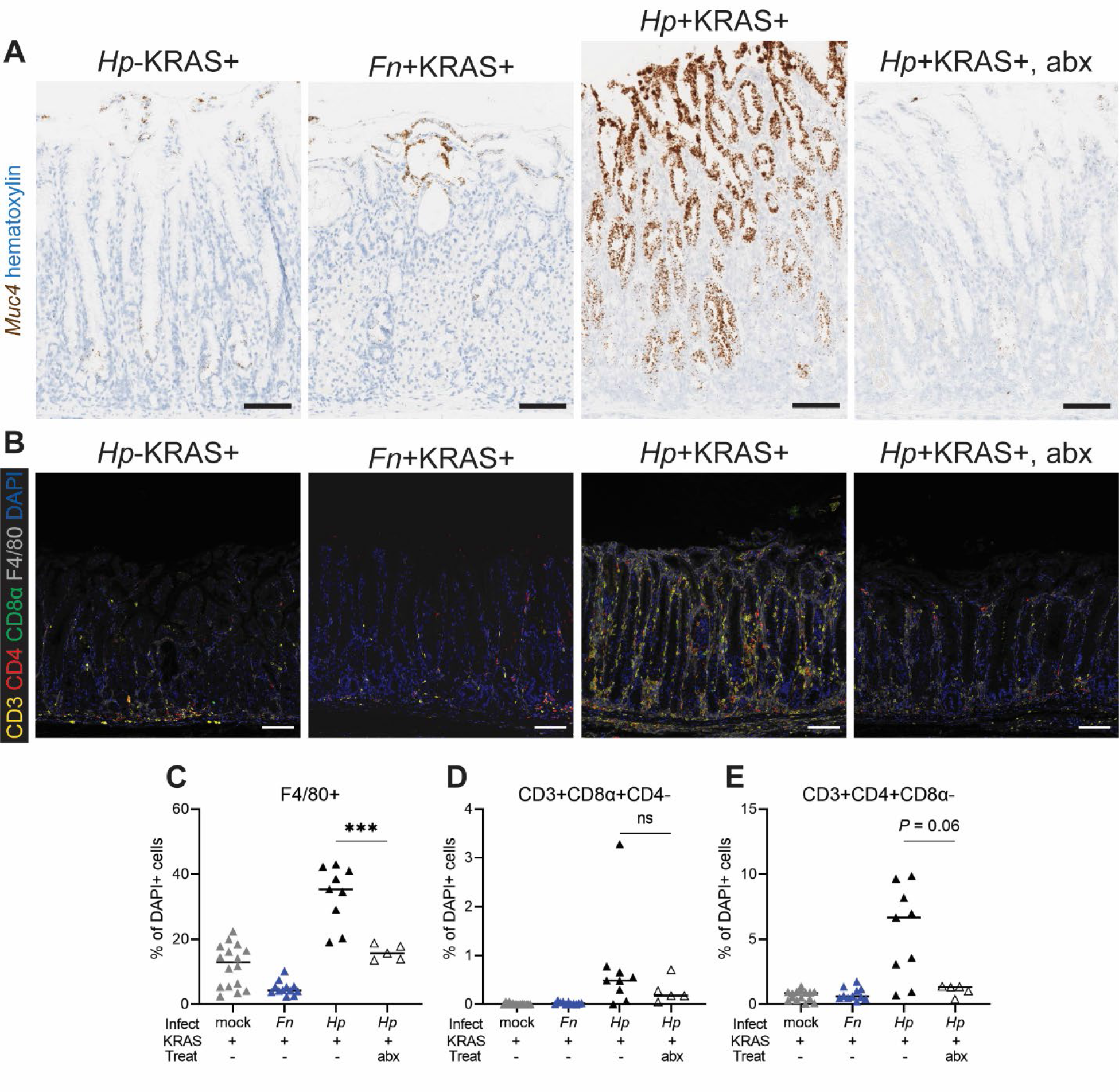
Metaplastic pit cell development requires sustained, *Hp*-specific inflammation. *Muc4* and gastric inflammation was assessed 12 weeks after KRAS induction in the following groups: *Hp*-KRAS+ mice; KRAS+ mice infected with *F. nucleatum* at six weeks (*Fn*+KRAS+); *Hp*+KRAS+ mice; and *Hp*+KRAS+ mice treated with antibiotics (‘abx’) from weeks six to eight. ISH was used to detect *Muc4* (brown) in the corpus at 12 weeks and representative images are shown. Scale bars, 100 µm. **B-E)** Immunohistochemistry (IHC) was used to detect the indicated immune cell subsets in gastric corpus tissue at 12 weeks. Data are from N=2 independent experiments with n=5-10 mice per group. **B)** Representative images of the gastric corpus are shown. CD3 (yellow) indicates T cells, CD4 (red) indicates helper T cells, CD8α (green) indicates cytotoxic T cells, F4/80 (grey) indicates macrophages and DAPI (blue) indicates nuclei. Scale bars, 100 µm. **C-E)** The indicated immune cell subsets were detected by IHC and quantified in each mouse using HALO software in N=2 experiments. Each datapoint is the average quantitation in two fields of view for an individual mouse and bars represent the medians. *** *P* < 0.001, Mann Whitney U test. ns, not significant.

One drawback to flow cytometry is the inability to visualize immune cell localization and distribution *in situ*. To further characterize gastric immunity in each mouse group, we used multiplex immunohistochemistry (IHC) (**Figure 4B**). Since our flow cytometry data (**Figure 3**) associated metaplastic pit cell expansion with an increased accumulation of macrophages and effector T cells in *Hp*+Kras+ mice, we focused on these immune cell subtypes. Each of these cell types was less abundant in *Hp*-KRAS+ and *Fn*+KRAS+ mice compared to *Hp*+KRAS+ mice. Quantitative tissue analysis using HALO software revealed that antibiotic treatment of *Hp*+KRAS+ mice caused a reduction in macrophages (F4/80+, *P* < 0.001, **Figure 4C**) and helper T cells (CD3+ CD4+CD8α-, *P* = 0.06, **Figure 4D**), but cytotoxic T cells (CD3+CD8α+CD4-) were not significantly impacted by antibiotic treatment (*P* = 0.30, **Figure 4E**). We observed that macrophages were localized throughout the lamina propria in all mouse groups. However, T cells were located throughout the lamina propria only in *Hp*+KRAS+ mice, whereas they were primarily located at the base of the gastric glands in the other treatment groups (**Figure 4B**), suggesting that T cell inflammation may impact a broader range of epithelial cell types in *Hp*+KRAS+ mice. Thus, metaplastic pit cell expansion requires sustained, *Hp*-specific gastric infection and associated inflammation.

### Metaplastic pit cells express amphiregulin and are not exclusively derived from transgene-expressing cells

We next explored whether *Muc4* expression was associated with other genes that could contribute to the tissue changes observed in *Hp*+KRAS+ mice. We determined the genes most strongly correlated with *Muc4* expression in metaplastic pit cells (**Figure 5A**) and found that one of the most strongly positively associated genes was *Areg* (amphiregulin), an epidermal growth factor receptor (EGFR) ligand that promotes epithelial cell growth and wound healing. We reasoned that amphiregulin could contribute to metaplastic pit cell expansion and the striking changes to gastric gland architecture seen in *Hp*+KRAS+ mice. We first assessed *Muc4* and *Areg* co-expression in each cluster in UMAP #2. Five clusters had a Pearson correlation *P* value < 0.05 indicating significant co-expression (**Figure 5B**). Expression of each marker was highest in pit_8 cells followed by pit_2 cells. Some neck_2 cells (which are likely SPEM cells, **Figure 2A**) as well as a few pit_3 and pit_4 cells also co-expressed these genes. Spatial transcriptomics revealed that the proportion of tissue spots that co-expressed *Muc4* and *Areg* was: 3.3% in *Hp*-KRAS-, 1.4% in *Hp*+KRAS-, 7.5% in *Hp*-KRAS+ and 17.8% in the *Hp*+KRAS+ sample (**Figure S6**). Finally, fluorescent ISH confirmed that *Muc4* and *Areg* strongly co- localized in *Hp*+KRAS+ mice at 12 weeks (**Figure 5C**). Because AREG does not trigger EGFR receptor internalization upon binding, downstream signaling pathways can be chronically activated ^27^. We performed Western blots on mouse stomach lysates at 12 weeks to query if metaplastic pit cell expansion correlated with EGFR protein levels. While there was no significant difference in EGFR levels between *Hp*-KRAS+ and *Hp*+KRAS+ mice, we detected a three-fold increase in phospho-EGFR in a *Hp*+KRAS+ mice (**Figure 5D**). These results suggest that the significantly increased *Areg* expression in these mice led to increased EGFR pathway activation.

**Figure 5.**
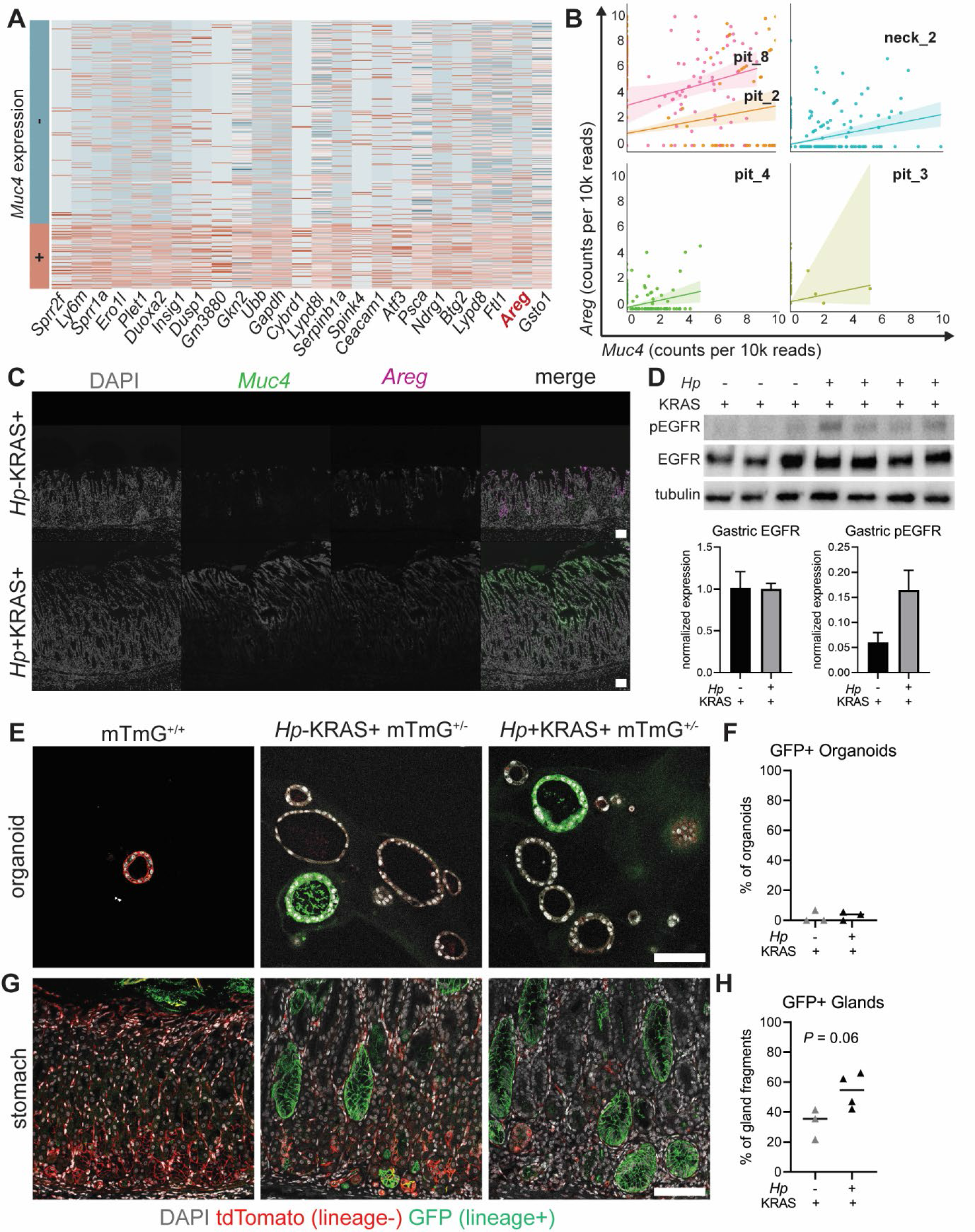
Metaplastic pit cells are associated with epidermal growth factor receptor signaling, but not with enhanced self-renewal. A) The heat map shows the normalized expression of the top 25 genes that are most significantly differentially expressed between *Muc4*-expressing (bottom) and non-expressing (top) metaplastic pit cells (pit_2, pit_6 and pit_8 cells). Genes are ranked from left to right in order of statistical significance. **B)** *Muc4* and *Areg* co-expression was assessed for each cluster in UMAP #2. Only the five indicated clusters had a Pearson correlation *P* value < 0.05. The expression of *Muc4* and *Areg* in counts per 10,000 reads is plotted for each cell in the given cluster. Trend lines are plotted and shading indicates the confidence interval of the linear estimation. **C)** Fluorescent ISH was used to detect *Muc4* (green) and *Areg* (cyan) from representative mice at 12 weeks; single channel images are shown in grayscale. Scale bars, 100 µm. **D)** EGFR and phosphorylated EGFR were detected in gastric cell lysates by Western blotting in from n=3 *Hp*-KRAS+ and n=4 *Hp*+KRAS+ mice from N=2 independent mouse experiments. Protein expression was normalized to tubulin and quantified by densitometry. **E-H)** Lineage tracing mice (*Mist1-CreERT2 Tg/+, LSL-Kras (G12D) Tg/+,* mTmG^+/-^) were infected with *Hp* or mock-infected and tamoxifen was used to induce constitutively active KRAS and GFP expression in *Mist1*-expressing cells. After 12 weeks, mice were euthanized and tdTomato and GFP expression was assessed in gastric organoids and tissues by immunohistochemistry. Untreated mTmG^+/-^ mice were included as controls. Data come from N=1 lineage tracing experiment with n=3-4 mice per group. Scale bars, 100 µm. **E-F)** Organoids were generated from the indicated mice and marker expression was assessed after 1-2 passages. **G-H)** Lineage-derived gland fragments were assessed in stomach tissues and quantified from 4-5 fields of view per mouse. The total percentage of GFP+ gland fragments is given; statistical significance was assessed with a Mann-Whitney U test. For **E** and **G**, individual channels are shown in Supplemental Figures S11 and S12.

To further explore changes to the gastric epithelium that occurred in *Hp*+KRAS+ mice, we performed a lineage tracing experiment using mTmG reporter mice, in which most cells express floxed tdTomato (red) at baseline. Upon tamoxifen treatment, Cre activity within a cell excises the tdTomato, allowing the expression of a downstream GFP (green) cassette. Thus, cells that express Cre, and all of their daughter cells, are GFP+ ^28^. We crossed our transgenic *Mist1-Kras* mice with mTmG mice, infected the progeny with *Hp* or mock-infected them, then administered tamoxifen to induce both Cre-mediated GFP expression for lineage tracing and constitutively active KRAS expression in chief cells. After 12 weeks, we generated organoids from the gastric corpus of three *Hp*-KRAS+mTmG^+/-^ and three *Hp*+KRAS+mTmG^+/-^ mice, cultured them under stem cell-promoting conditions for 1-2 passages, and then immunostained them for tdTomato and GFP (**Figure 5E, S11**). Gastric organoids are derived from gastric epithelial stem cells, which have the unique property of self-renewal, a process by which stem cells divide to generate more stem cells ^29^. We observed that only a low proportion of organoids were GFP+, regardless of whether the organoids were generated from mice with *Hp* infection (0-7 GFP+% in *Hp*-KRAS+mTmG^+/-^ and 0-6% in *Hp*+KRAS+mTmG^+/-^ mice, **Figure 5F**). The overall low proportion of GFP+ organoids suggested that whether KRAS+ mice had *Hp* infection or not, a similar proportion of lineage-derived cells exhibited the stem cell property of self-renewal.

Intriguingly, we observed that the organoids from KRAS+ mice that were GFP- were not tdTomato+, which could suggest that they did not express the mTmG reporter construct. Control organoids from an mTmG mouse were uniformly tdTomato+ (**Figure 5E**). To explore gastric epithelial expression of the mTmG construct, we immunostained thin sections of gastric corpus tissue for tdTomato and GFP. In control mTmG mice that did not receive tamoxifen, expression of tdTomato was most evident in cells at the base of the glands (e.g., chief cells), suggesting that other epithelial cells (e.g., neck, parietal, pit and isthmal stem cells) express lower levels of the mTmG construct, in spite of its presence in all cells. In these control mice there was no GFP expression because these mice did not have activation of Cre (**Figure 5G**, left, note the green signal at the top of the glands is autofluorescence of stomach contents, e.g., food). In *Hp*- KRAS+mTmG^+/-^ and *Hp*+KRAS+mTmG^+/-^ mice at 12 weeks (the same mice from which organoids were generated), tdTomato+ glands were scant (**Figure 5G, S12**). However, we observed more GFP+ glands in *Hp*+KRAS+ mice than in *Hp*-KRAS+ mice (**Figure 5H**): the median proportion of GFP+ glands was 36% in *Hp*-KRAS+ mice and 55% in *Hp*+KRAS+ mice, demonstrating that *Hp* infection sustains and/or expands lineage-positive glands in this model. Importantly, by ISH we observed expression of *Muc4*, our marker of metaplastic pit cells, in up to ∼100% of corpus glands in *Hp*+KRAS+ mice (see **Figure 2E, 4A**). Thus, up to half of the glands in *Hp*+KRAS+ mice may harbor metaplastic pit cells, yet are lineage-negative. These results suggest that metaplastic pit cells are not directly derived from cells in which the *Kras* transgene was induced; instead, the combination of *Hp* infection-associated inflammation plus induction of constitutively active KRAS generates a gastric microenvironment that favors metaplastic pit cell expansion. However, the organoid experiments demonstrate that this unique gastric environment does not skew the balance of lineage-positive cells that exhibit self-renewal. Thus, the cell state changes in *Hp*+KRAS+ mice likely fall short of cancer ^30^.

### MUC4 is associated with cell proliferation in subjects with gastric cancer

Gastric cancers often consist of mixed cell types including pit or pit-like cells ^31, 32^, but the significance of pit cells in disease development is not well understood. Thus, we used a tissue microarray (TMA) to assess mucin expression in samples from 47 recent gastric cancer cases from the United States Pacific Northwest (**Table S6**). Samples consisted of two-millimeter tissue cores from regions of superficial cancer, deep cancer and non-neoplastic epithelium adjacent to cancer. One section was immunostained for the classical pit cell mucin MUC5Ac and the intestinal mucin MUC2 (**Figure 6A**), and a second section was immunostained for MUC4 and the proliferation marker Ki-67 (**Figure 6B**). We used QuPath to segment cells via nuclear staining and quantify marker expression based on fluorescence intensity ^33^ (**Table S7**). Marker expression differed according to tissue type (**Figure 6C**). Expression of the pit cell mucin MUC5Ac was higher in the non-neoplastic epithelium and superficial cancer than in deep cancer. Expression of the intestinal MUC2 was lower in non-neoplastic epithelium than in either cancer type. Both MUC4 and the proliferation marker Ki-67 had the highest expression in superficial cancer compared to the other tissues; as well, Ki-67 expression in deep cancer was higher than in non-neoplastic epithelium.

**Figure 6.**
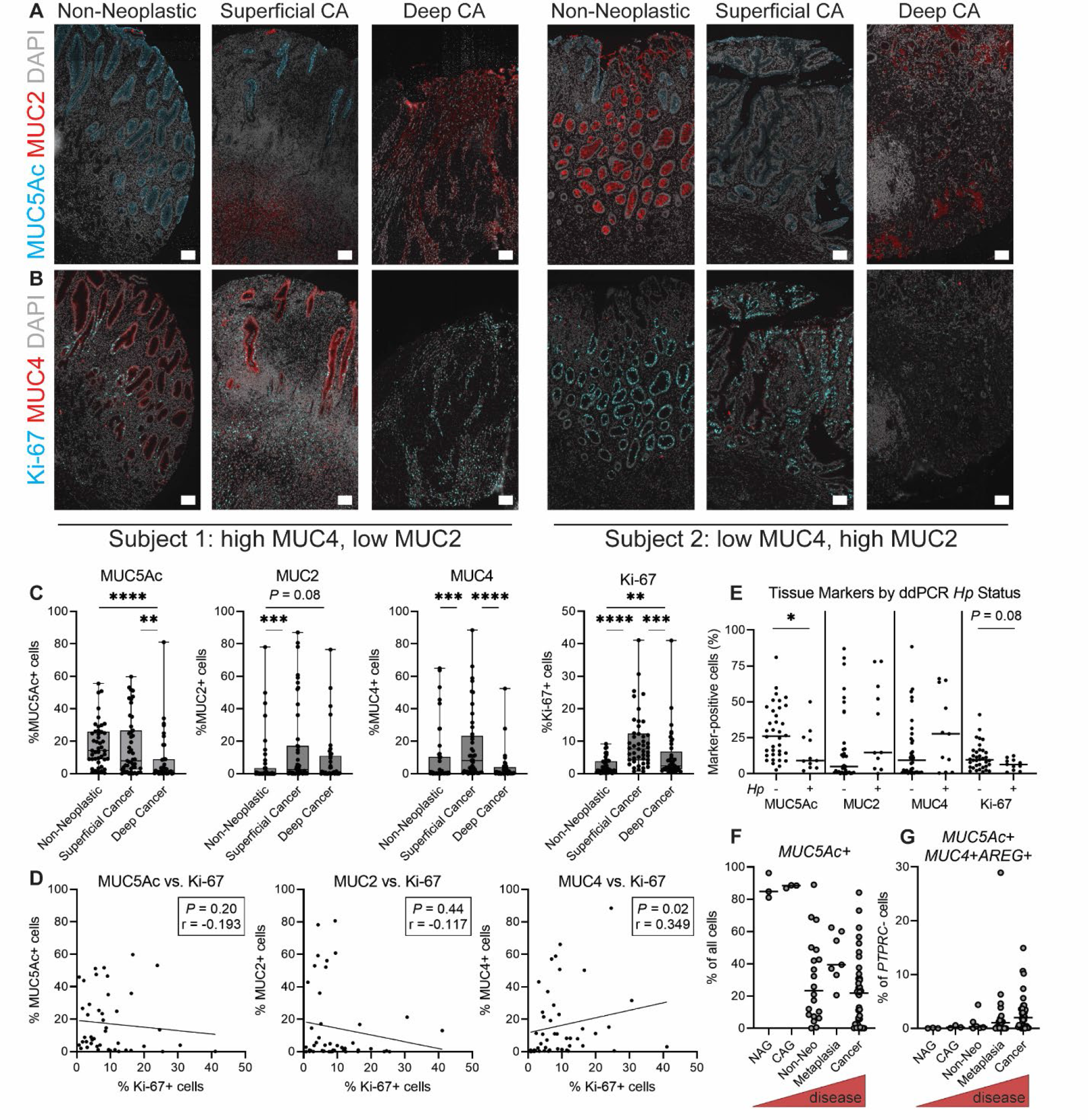
Metaplastic pit cells are detected in human subjects with preneoplasia and gastric cancer. A-E) The expression of the mucins MUC5Ac, MUC2 and MUC4 and the cell proliferation marker Ki-67 was probed in a tissue microarray (TMA) comprising samples from 47 gastric cancer (“CA”) patients. **A-B)** Marker expression is shown in tissue sections from two representative patients. Scale bars, 100 µm. **A)** MUC2 is shown in red, MUC5Ac in cyan and DAPI in grey. **B)** MUC4 is shown in red, KI-67 in cyan and nuclei (DAPI) in grey. **C-D)** For each tissue core, QuPath was used to segment individual cells and determine their marker expression by pixel intensity. **C)** The percentage of positive cells was calculated for each marker and sample, and is shown as box-and-whisker plots overlaid with individual values for each sample. **D)** The Spearman correlation between the percent of Ki-67+ cells and the percent of mucin+ cells is given for each superficial cancer sample. **E)** Gastric *Hp* status was determined by droplet digital PCR and subjects were categorized as *Hp*- or *Hp*+ to indicate whether their stomachs still harbored active *Hp* infection. The percentage of positive cells is shown for each marker and subject according to the subjects’ *Hp* status. The tissue core with the greatest expression of each marker was used. **F-G)** Three published gastric scRNA-seq datasets were mined for cell types of interest. Each dot represents one scRNA-seq sample, which are arranged in order of disease severity. **F)** Pit cells (*MUC5Ac*+) are shown as the proportion of all cells in each sample. **G)** Metaplastic pit cells (*MUC5Ac*+*MUC4*+*AREG*+) are shown as the proportion of non-hematopoietic cells (*PTPRC*-) in each sample. NAG, non-atrophic gastritis; CAG, chronic atrophic gastritis. In **C** and **E** significance was calculated using Mann-Whitney U tests; * *P* < 0.05, ** *P* < 0.01, *** *P* < 0.001, **** *P* < 0.0001. In **D** significance was calculated using Spearman correlations and the *P* values and correlation coefficients (r) are given; the trend lines were fit using simple linear regressions. In **E, F** and **G** bars indicate median values.

We next investigated correlations between the different markers within each subject, focusing on the superficial cancer samples because these tissues had relatively high expression of each marker. Only MUC4 was significantly correlated with Ki-67 expression (Spearman *P* < 0.05), with a modest positive association (r = 0.349, **Figure 6D**). MUC5Ac and MUC2 were each negatively associated with Ki-67, though the correlations were not statistically significant.

Because MUC4 and Ki-67 were included in the same staining panel, we probed their expression in each individual cell from all tissue cores, for a total of 2.69 million cells. MUC4 and Ki-67 were again positively correlated (**Figure S13**). Finally, we asked whether marker expression was correlated with active *Hp* infection. We used droplet digital PCR to detect *Hp* in DNA extracted from the tissues used to generate the TMA. Eleven of 47 subjects had detectable *Hp* DNA, consistent with previous observations that at least half of gastric cancer patients clear *Hp* infection prior to cancer diagnosis ^34–36^. Due to the heterogeneity in marker expression among sample types, we analyzed the tissue type with the greatest marker expression for each subject. *Hp*- subjects had more MUC5Ac+ and Ki-67+ cells than *Hp*+ subjects (**Figure 6E**). The median level of MUC2 and MUC4 expression was higher in *Hp*+ subjects than *Hp*- subjects, though the differences were not statistically significant (**Figure 6E**). Because of substantial heterogeneity in marker expression, we analyzed the same data to determine the proportion of subjects with high vs. low expression of each marker (**Figure S14A**). *Hp*+ subjects were more likely to have high MUC2 and MUC4 expression, whereas *Hp*- subjects were more likely to have high MUC5Ac and Ki-67 expression.

Finally, to explore how pit cells change over the course of human disease, we mined data from three recently published gastric scRNA-seq studies (**Figure 6F-G**). One study had nine subjects with 13 antral biopsies: three with non-atrophic gastritis (NAG, mild gastric inflammation), three with chronic atrophic gastritis (CAG, loss of parietal and chief cells due to *Hp*-mediated inflammation), six with metaplasia and one with cancer ^37^. The other two studies comprised 20 samples from seven subjects with cancer and one with metaplasia ^38^ and 48 samples from 31 subjects with cancer ^39^; these studies also included paired samples from non-neoplastic epithelium adjacent to cancer. We observed that *MUC5Ac*+ cells, presumably pit cells, comprised a large proportion of cells sequenced in NAG and CAG samples, i.e., mild disease. As disease severity increased, the proportion of *MUC5Ac+* (pit) cells detected from each sample decreased (**Figure 6F**). We found that cells expressing *MUC5Ac*, *MUC4* and *AREG*, i.e., metaplastic pit cells, made up an increasing proportion of non-hematopoietic cells (*PTPRC-*) as disease progressed (**Figure 6G**). Among the samples from later stages of disease, these cells comprised 0.03-4.4% (median 0.29%) of non-hematopoietic cells in samples of metaplasia, 0-28.9% (median 1.1%) of non-hematopoietic cells in non-neoplastic tissue adjacent to cancer, and 0-14.9% (median 2.0%) of non-hemopoietic cells in cancer samples. As well, as disease progressed, pit cells had greater expression of *MUC2* and *MKI67* (Ki-67, **Figure S14B**). Finally, we confirmed in our TMA that in human subjects with gastric cancer, pit cells (expressing the MUC5Ac transcript or protein) could express the *MUC4* transcript or MUC2 protein (**Figure S15**). Thus, pit cell expression of the intestinal mucins MUC2 and MUC4 coincides with gastric disease severity

## Discussion

The common hypothesis for how *Hp* infection leads to cancer is that *Hp*-driven inflammation causes the accumulation of oncogenic mutations and activation of oncogenic pathways, which can lead to metaplasia and cancer development even if the host clears *Hp* infection. In other words, *Hp* is essential for <u>initiating</u> mutagenic gastric inflammation, but once oncogenic pathways become activated, *Hp* is dispensable. However, up to half of gastric tumors still harbor active *Hp* infection ^34–36^, and eradication of *Hp* in combination with surgical tumor resection significantly reduces cancer recurrence, compared to surgical resection without *Hp* eradication^15^. Additionally, eradication of *Hp* in subjects with metaplasia significantly reduces their risk of developing gastric cancer ^13, 14^. Cumulatively, these observations suggest that active *Hp* infection during the later stages of disease promotes cancer development. Here we used a mouse model of gastric metaplasia to test how the additional perturbation of chronic *Hp* infection impacts disease. We found that the combination of *Hp* infection and metaplasia driven by induction of a constitutively active *Kras* allele (*Hp*+KRAS+ mice) elicited a striking expansion of metaplastic cells most similar to surface mucus-producing pit cells. Metaplastic pit cells expressed the intestinal mucin *Muc4* and the EGFR ligand amphiregulin and were dependent on *Hp*-specific inflammation. Thus, metaplasia in *Hp*+KRAS+ mice is not simply accelerated beyond what is seen in *Hp*-KRAS+ mice, but instead develops along an altered trajectory. We detected *Muc4* expression in our human gastric cancer subjects, especially in the context of current *Hp* infection, but organoid studies revealed that lineage-traced cells from *Hp*+KRAS+ mice had limited capacity for self-renewal. Thus, additional genetic or epigenetic changes are likely needed to drive *Hp*-associated metaplastic pit cells to cancer. These data support the hypothesis that *Hp* can actively impact gastric metaplasia, beyond simply initiating inflammation.

Gastric tumors often harbor pit or pit-like cells ^31, 32^ but the role of pit cells in preneoplasia and tumorigenesis is not well understood. A meta-analysis found that during gastric cancer, decreased expression of the pit cell mucin MUC5Ac was associated with deeper tumor penetration and worse overall survival ^40^. Here we found that *MUC5Ac*+ cells decreased in overall abundance as disease progressed in three scRNA-seq studies using human samples.

However, as pit cells decreased in abundance, they began to express the intestinal mucins *MUC2* and *MUC4*. *MUC5Ac*+ cells have been shown to express *MUC2* in some patients with intestinal metaplasia ^41^. Previous studies observed MUC4 in human gastric tumors but did not examine cell type-specific expression ^42, 43^. As a transmembrane mucin, MUC4’s extracellular domain can interact with the extracellular environment (which could include interactions with pathogens such as *Hp*), while its cytoplasmic domain can promote oncogenic signaling through multiple pathways, including HER2 (Erbb2) interactions and activation of Notch3 signaling ^44^.

Our gene set enrichment analysis implicated ErbB signaling, which could involve both HER2 and ErbB1 (EGFR), and we found that the expression of *Muc4* was highly correlated with the expression of the EGFR ligand amphiregulin. Most EGFR ligands (e.g., EGF, TGFβ) bind with high affinity to EGFR and induce EGFR receptor internalization, transiently impairing EGFR function and driving oscillation of MAP kinase signaling. In contrast, amphiregulin (AREG), a low-affinity ligand, does not drive EGFR internalization, and instead leads to tonic signaling that can promote both cell differentiation and proliferation ^27^. The observation that *Hp*+KRAS+ mice had increased levels of phospho-EGFR in stomach lysates supports a model where *Areg* expression in metaplastic pit cells may promote or sustain their expansion. As well, because amphiregulin can function in autocrine, paracrine and juxtacrine fashion ^45^, induction of *Areg* in metaplastic pit cells could leads to widespread changes in the stomach through paracrine signaling. Interestingly, *Areg* knockout mice were reported to spontaneously develop antral tumors over the course of 18 months ^46^, suggesting that amphiregulin protects against gastric cancer. However, *AREG* upregulation has been reported in gastric cancers as well ^47, 48^. These observations underscore the importance of amphiregulin in stomach physiology and imply that precise regulation of *Areg* is critical to prevent disease. As there are now several therapies available to target EGFR signaling during cancer, more work is needed to clarify the role(s) of amphiregulin in our model and in gastric cancer.

We found that MUC4 was significantly positively associated with the cell proliferation marker Ki-67 in individual cells from human subjects with gastric cancer. However, organoids generated from *Hp*+KRAS+ mice were no more likely to be lineage-derived than organoids from *Hp*-KRAS+ mice, suggesting that metaplastic pit cells have limited capacity for self-renewal *ex vivo*. As well, *Hp*+KRAS+ mice did not develop gastric tumors within a three-to-four month experimental time frame ^16^. Thus, metaplastic pit cells may serve as a precursor cell type that is poised to become cancerous in certain contexts. In our model, KRAS G12D is induced in *Mist1*- expressing cells, which are primarily chief cells but also a quiescent isthmal stem cell population ^49^. We cannot rule out a role for *Mist1*-expressing isthmal stem cell expansion in *Hp*+KRAS+ mice. However, pit cell hyperplasia has been seen in other KRAS G12D models, including *Lrig1-Kras* mice ^50^, *eR1-Kras* mice (targeting Runx1) ^51^ and *Tff1-Kras* mice ^52^, in the absence of *Hp* infection. Interestingly, *Tff1-Kras* mice had robust MUC4 expression in the gastric epithelium and increased expression of *Areg* compared to healthy control mice. Thus, multiple gastric perturbations can lead to MUC4 and amphiregulin expression, suggesting that this may be a conserved response to gastric injury.

Previously, lineage tracing of chief cells that transdifferentiated due to pharmacologically induced parietal cell loss showed that almost 25% of lineage-positive cells expressed MUC5Ac, suggesting that SPEM cells can convert to pit or pit-like cells ^5^. However, the majority of pit cells were lineage-negative, suggesting that while SPEM cells may contribute to foveolar (pit cell) hyperplasia, they are not the primary driver. *Hp* infection is another perturbation known to promote foveolar hyperplasia as well as changes in pit cell gene expression and phenotypes ^53, 54^. Thus, the combination of *Hp* infection plus SPEM due to transgene induction in chief cells may act synergistically to drive pit cell metaplasia in our model. Whether the two perturbations increase SPEM cell conversion into pit-like cells, or may generate an inflammatory gastric milieu that skews pit cells or isthmal progenitors toward metaplasia, is not yet known, in part due to the limitations of our lineage tracing studies. Intriguingly, laser-capture microdissection of pit cells from *Hp*-infected Balb/c mice showed a signature of IFNγ response and actin rearrangement ^53^, two key responses that were upregulated in metaplastic pit cells compared to pit cells. The cytokine IFNγ has been shown to induce MUC4 expression in cultured cell lines, both individually ^55^ and synergistically with TNFα ^56^. These cytokines were also implicated by our gene set enrichment analysis comparing metaplastic versus classical pit cells. Our finding that dexamethasone treatment prevented metaplastic pit cell expansion supports a role for *Hp*- mediated inflammation in driving this cell type. In this study we focused on macrophages and T cells, not only because these immune cell populations were significantly correlated with metaplastic pit cell development but also because these cells have been associated with gastric disease in other animal models ^4, 12, 24, 57^. However, dexamethasone is broadly immunosuppressive and we cannot rule out the contribution of other immune cell types.

Intriguingly, long-term use of non-steroidal anti-inflammatory drugs (NSAIDs) is associated with reduced risk of noncardia gastric cancer ^58^.

Our study has several limitations. i) It may be that the specific changes in disease marker expression observed in *Hp*+KRAS+ mice are due to effects on *Mist1*-expressing isthmal progenitors ^49^ in addition to, or instead of, effects on chief cells. However, a study that used the highly chief cell-specific reporter GIF (gastric intrinsic factor) confirmed that chief cells are the primary cell of origin of SPEM ^5^. ii) Our lineage analysis was limited by low expression of the tdTomato reporter in many cell types. However, we did detect more GFP+ glands in *Hp*+KRAS+ mice than in *Hp*-KRAS+ mice, suggesting that the unique gastric environment in *Hp*+KRAS+ mice may either sustain or expand lineage-derived cells. We note that EGFR signaling induced by *Areg* expression in metaplastic pit cells could lead to RTK-RAS signaling, which would preclude the need for KRAS G12D induction within a given cell type. iii) Gastric cancer is quite heterogeneous and RAS pathway activity is only found in ∼40% of tumors ^59^. However, we and others previously showed that induction of constitutively active KRAS in *Mist1*-expressing cells is sufficient to drive metaplasia and dysplasia, thus establishing that constitutively active KRAS is a tool to broadly model aspects of human disease. iv) We observed that human subjects with constitutively active *Hp* infection detected by ddPCR were more likely to have gastric MUC2 and MUC4 expression. However, only 11 of 47 subjects had evidence of *Hp* by ddPCR, limiting the statistical power of these analyses. We note that about 80% of gastric cancers are attributed to *Hp* infection, so it is likely that the majority of our subjects had an *Hp* infection that was either cleared prior to cancer diagnosis or was below our limit of detection. In a previous study with 63% *Hp* positivity, *Hp* was significantly positively associated with both MUC2 and MUC4 expression in gastric tumors ^60^. v) Finally, we note that although *Fusobacterium nucleatum* did not drive metaplastic pit cell development, the bacterium did not colonize the stomach as robustly as *Hp* did. However, in *Hp*+KRAS+ mice, *Muc4* levels were not associated with *Hp* loads, so the lack of *Muc4*expression in *Fn*+KRAS+ mice is likely due to their reduced inflammation (which could result from overall lower bacterial loads).

Our mouse model provides a means to dissect the cell lineages that arise during infection-associated preneoplastic progression. In the present study we discovered that *Hp* infection during gastric metaplasia leads to the expansion of *Muc4*-expressing metaplastic pit cells. These cells were highly expanded in *Hp*+KRAS+ mice as evidenced by the greatly increased abundance of the pit_2, pit_6 and pit_8 clusters and the widespread *Muc4* detected by *in situ* hybridization. In published single sequencing data from human gastric cancer and precancer subjects, *MUC4* expression was greatest in pit cells from subjects with metaplasia and cancer. In our mouse model, lineage-derived cells in the *Hp*+KRAS+ stomach had limited capacity for self-renewal. Thus, we propose that the metaplastic pit cell phenotype represents *Hp*-driven gastric metaplasia rather than cancer. More broadly, we note that most patients with metaplasia do not progress to cancer, but MUC4 expression was strongly linked with Ki-67 expression in gastric cancer tissues in our human cohort. Future studies will test whether the presence of metaplastic pit cells, especially MUC4-expressing pit cells, during gastric metaplasia may serve as a risk factor for progression from metaplasia to cancer.

## Supplemental Methods

### Data availability statement

Single-cell RNA-sequencing data is being deposited in GEO and will be publicly available on January 31, 2023. The accession numbers are: SAMN31671449, SAMN31671450, SAMN31671451, SAMN31671452, SAMN31671453, SAMN31671454, SAMN31671455, SAMN31671456.

### Mouse model of gastric preneoplasia

All mouse experiments were approved by the Fred Hutchinson Cancer Center Institutional Animal Care and Use Committee (protocol number 1531) and were performed in accordance with the recommendations in the National Institutes of Health Guide for the Care and Use of Laboratory Animals. *Mist1-CreERT2 Tg/+, LSL-Kras (G12D) Tg/+* (“*Mist1-Kras*”) mice were previously described ^4, 16^. Male and female mice aged eight to 16 weeks old were randomized to treatment groups. On day one, mice were infected with *Hp* strain PMSS1 ^61^ or mock-infected with broth. On days two through four, mice were given 5 mg tamoxifen (Sigma) in corn oil (Sigma) by subcutaneous injection, or were given corn oil alone. Mice were humanely euthanized after two, six or 12 weeks and tissues were analyzed. For lineage tracing studies, *Mist1-Kras* mice were crossed with “mTmG” reporter mice ^28^ (membrane-localized mTmG, *Gt(ROSA)26Sortm4(ACTB-tdTomato,-EGFP)Luo*/J, #007576, the Jackson Laboratory).

### Bacterial infections

*Helicobacter pylori* strain PMSS1 was cultured and mice were inoculated with 10^8^ CFU as previously described ^16^. *Fusobacterium nucleatum* subsp. *animalis* strain COCA36 ^25^ was streaked onto Fastidious Anaerobe Agar plates (46 g agar per liter water, Grainger) containing 10% defibrinated horse blood (HemoStat) and JVN antibiotics (3 µg/ml josamycin, Fisher; 4 µg/ml vancomycin hydrochloride, Fisher; and 1 µg/ml norfloxacin, Fisher) and cultured at 37°C in an anaerobic chamber (Coy). After two days of growth on antibiotics, colonies were expanded onto 10% horse blood plates without antibiotics for two additional days. An inoculum of 10^8^ CFU was prepared by scraping bacteria from the plates and resuspending in phosphate-buffered saline. To determine *Fn* stomach titers, harvested tissues were weighed, homogenized, serially diluted and plated on 10% horse blood with JVN antibiotics in an anaerobic chamber. “Triple therapy” antibiotic treatments were previously described ^16^. For immunosuppression with oral corticosteroids, because the Fred Hutch small animal vivarium acidifies its water to pH ∼3.5, all mice were given autoclaved, non-acidified water starting at two weeks after infection and constitutively active KRAS induction. Dexamethasone (Sigma) was added to the drinking water at 1 mg/liter. Control animals received non-acidified water only. Water was protected from light and changed weekly and mice were euthanized at six weeks.

### scRNA-seq of the gastric corpus

Stomachs were aseptically harvested. One third was removed and fixed in 10% neutral-buffered formalin. The remainder of the tissue was trimmed to remove the forestomach and antrum and transferred to ice-cold PBS. Tissues were washed by agitating in three serial passages of ice-cold PBS, then single-cell suspensions were generated using digestion at 4°C with protease from *Bacillus licheniformis* (Sigma P5380). An established protocol was used (see ^62^ section 3.2 – Adult Mouse Lung) with the following modifications: instead of weighing 25 mg of tissue, stomach tissue was minced and then divided in half, and each half was used in a protease digestion reaction; samples were rocked gently on a rocker in between trituration steps; and a cell strainer was not used as straining led to epithelial cell loss. After single cell suspensions were generated, some samples were subjected to cryopreservation. Briefly, cells were stored in Recovery Cell Culture Freezing Medium (Gibco), cooled slowly overnight in a CoolCell Container (BioCision) placed at -80°C, then stored long-term in liquid nitrogen. Cells were rapidly thawed in a 37°C water bath. At the 6 week time point, cells were thawed from one *Hp*- KRAS+ mouse and one *Hp*+KRAS+ mouse. At the 12 week time point, thawed cells were pooled from 2-3 mice per treatment group (described in **Table S1**). Dead cells were removed using magnetic bead separation (MACS Dead Cell Removal Kit with MS columns, Miltenyi Biotec). Live cells were pelleted, resuspended in phosphate-buffered saline with 0.04% non-acetylated bovine serum albumin, and counted with a hemacytometer using Trypan Blue (0.4%, Gibco) to distinguish between live (clear) and dead (blue) cells.

In an attempt to increase the number of sequenced cells, two samples from the 12 week time point were not subjected to cryopreservation. Instead, the single cell suspensions were generated from one *Hp*-KRAS+ and one *Hp*+KRAS+ mouse using cold protease digestion as described above, then dead cells were immediately removed using magnetic bead separation as described above. The live cells were then counted and immediately used.

Whether samples were cryopreserved or not, after dead cell clean up, cells were used for gel beads-in-emulsion (GEM) generation and barcoding using the Chromium Next GEM Single Cell 3ʹ Reagent Kits v3.1 and Nextera library preparation (both from 10x Genomics) according to manufacturer’s instructions. After standard library quality control metrics, samples were sequenced in a HiSeq rapid flow cell with an Illumina HiSeq 2500 sequencer (six week time point) and a NovaSeq SP 100 flow cell with an Illumina NovaSeq 6000 sequencer (12 week time point).

### scRNA-seq analysis

#### Quality control and dimension reduction

Sequencing reads were aligned to the mouse genome using cellRanger (10x Genomics). Seurat was applied to filter the output feature count matrixes of these samples to include only cells expressing at least 250 genes and genes expressed in at least 10 cells (18,089 genes and 22,050 cells passed this filter from a starting population of 6.7 million GEMs), and to filter out cells with >25% mitochondrial content (16,434 cells remained after this filter). Filtered cells were integrated into a single dataset using the default Seurat parameters. The integrated reads were normalized, scaled and subjected to principal component analysis (PCA) dimension reduction. A SNN clustering method was applied on the processed dataset to find 25 clusters using the first 20 dimensions after PCA reduction and resolution of 0.5. Clusters were then visualized on a 2- dimensional UMAP plot. We applied Seurat’s FindAllMarkers function to identify the representative marker genes for each cluster and assigned cell types accordingly. To explore the predominant gastric epithelial cell populations, cells from the central epithelial “megacluster” in the 12 week samples were separated and re-clustered into 14 clusters using the same Seurat processing workflow described above, with a resolution of 0.7. To generate forest plots and heat maps showing cell abundance, samples were categorized according to treatment and time (i.e., the two 12 week *Hp*-KRAS+ and the two 12 week *Hp*+KRAS+ samples were each condensed into one sample). For each group, the proportion and confidence interval of each cell type was estimated from the empirical Bayesian distribution based on the observations of cell type occurrence in R using the EBBR package. The corresponding forest plots and heat maps were generated using ggplot2. Forest plot error bars represent the confidence interval that a given cell would be identified as a given cell type based on the observed cell distributions in the dataset. Heatmaps showing gene expression in different cell types and conditions were generated in Python via the Pandas and Seaborn packages using the average value of normalized gene expression of all cells from a given cell type or condition. Contour plots for *Muc5ac* and *Muc4* expression were generated with Seaborn package using scaled and normalized gene expression data from Seurat.

#### Gene Set Enrichment Analysis

We applied the FindAllMarkers function from Seurat (minimum 10% cells expressing that gene, minimum log2 fold change of 0.25 between the two groups) to achieve a list of differentially expressed genes between clusters pit_2, pit_6 and pit_8 (metaplastic pit cells) vs. pit_1, pit_3, pit_4, pit_5 and pit_7. Statistically significant genes (false discovery rate [FDR] <= 0.05, Wilcoxon rank sum test) were used as the input for downstream GSEA (Gene Set Enrichment) analysis (GSEA Pre-ranked, fold change value as ranking order) using Hallmark, KEGG and GO pathway datasets from MSigDB ^63, 64^. A 0.15 FDR value was used as cutoff for statistically significantly enriched pathways.

#### Spatial gene expression (10x Visium)

Four mice were used: *Hp*-KRAS-, *Hp*+KRAS-, *Hp*-KRAS+ and *Hp*+KRAS+, each at the 12 week time point. Mouse stomachs were embedded in OCT and snap-frozen in isopentane/liquid nitrogen, then stored at -80°C. Using a cryotome, sections of 10 µm thickness were cut onto a Visium slide (10x, serial number V11F01-280) with two sections (from the same mouse) per capture area. Adjacent sections were cut onto glass slides and stained with Diff-Quick (Differential Quik III Stain Kit, Polysciences) to assess tissue morphology. On the day of the experiment, the slide was fixed and stained according to the manufacturer’s instructions. After washing, the slide was mounted in 85% glycerol (Fisher) with 2 U/µl Protector RNase inhibitor (Millipore Sigma) and imaged on a Leica Microsystems DMi8 scanning microscope. Tissue sections were permeabilized for six minutes and RNA was extracted and library prepped according to manufacturer’s instructions. The libraries were pooled with another Visium sample and sequenced on a NovaSeq SP 100 flow cell with an Illumina NovaSeq 6000 sequencer, then deconvoluted post-sequencing. The multicolor LIF and TIFF images were viewed in SpaceRanger (10x Genomics) and fiducial markings were used to align the Visium spots.

Regions of folded or damaged tissue were discarded. The cleaned data was log-normalized and PCA-reduced based on highly variable genes with BayesSpace ^65^ using the default settings.

The number of clusters for each sample was defined as the elbow point of the qPlot, using the first 15 dimensions of the PCA. Feature expression plots were generated using the BayesSpace-enhanced data via the enhanceFeature function from BayesSpace.

#### Analysis of published human scRNA-seq datasets

We collected three human scRNA datasets from subjects with gastric pre-cancer or early gastric cancer ^37–39^ to validate the existence of metaplastic pit cells in the human population. The feature count matrix from each subject was fed into the standardized Seurat processing pipeline. Sequencing data were filtered by Seurat default settings to remove cells with high mitochondrial mRNA amounts or low gene expression. Filtered data were normalized and scaled and the number of cells expressing a certain gene was determined by the subset function in R.

### Flow Cytometry

Stomachs were aseptically harvested, the antrum and forestomach were discarded, and the remaining corpus tissue was kept on ice in RP-3: RP-0 (RPMI medium [Gibco] containing 1% penicillin/streptomycin/L-glutamine [Gibco] and 1% HEPES [Cytiva]), with the addition of 3% HyClone fetal bovine serum (Cytiva). Tissues were opened, rinsed with a basic buffer (10 mM HEPES, pH 8.2) and cut into 3-4 pieces, which were incubated with 400 rpm stirring for 20 minutes at 37°C in 10 ml RP-3 with 5 mM EDTA (Sigma) and 0.15 g/L DTT (Sigma). Tissue pieces were vigorously shaken three times for 30 seconds each in 7 ml RP-0 with 0.5M EDTA and strained over a kitchen strainer to remove epithelial cells and intraepithelial lymphocytes. The tissue pieces were finely minced with scissors, then incubated with 400 rpm stirring for 25 minutes at 37°C in 5-7 ml of a digestion buffer of RP-0 with 10 µg/ml DNase I (Roche) and 0.2 mg/ml Liberase TL (Roche). The enzymatic reaction was stopped with the addition of 10 ml cold RP-3 and tissues were filtered through a 70 µm strainer. The remaining tissue pieces were gently mashed with the rubber end of a 1 ml syringe. Cells were pelleted and resuspended in 5 ml 37.5% Percoll (Cytiva), then centrifuged for 20 mins at room temperature in an Eppendorf benchtop centrifuge (5810 R) at 1800 rpm with Ascend and Descend set to 1. The supernatant was aspirated and the pellet, containing lamina propria immune cells, was resuspended in 270 µl RP-3, of which 100 µl was used to detect myeloid cells and 100 µl was used to detect lymphocytes as follows. Briefly, cells were incubated in 40 µl of 1:1000 Ghost Dye R780 fixable viability dye (Tonbo). This and all subsequent steps were performed in the dark at room temperature unless otherwise stated. After 20 minutes, 10 µl of 1:400 Fc block was added (anti-mouse CD16/32, 2.4G2, BD). After 10 minutes, 50 µl of the extracellular antibody stain was added (see **Table S8**). After 30 minutes, cells were pelleted. Cells stained with the myeloid panel were resuspended in 200 µl of 2% paraformaldehyde. Cells stained with the lymphocyte panel were resuspended in 200 µl of Fix/Perm (FoxP3/Transcription Factor Staining Buffer Set, eBioscience, diluted according to manufacturer’s instructions) and incubated for 20 minutes at 4°C. Cells were pelleted, washed in 200 µl PermWash (eBioscience, diluted according to manufacturer’s instructions) and incubated with 100 µl of intracellular antibody stain (see **Table S8**) in PermWash for 30 minutes. Cells were pelleted and resuspended in 200 µl FACS buffer (PBS with 2% fetal bovine serum and 1 mM EDTA). Finally, 50 µl of the 270 µl cell preparation was stained with 1:1000 Ghost Dye (R780) and fixed with 2% paraformaldehyde containing 20,000 counting beads (Accucheck Counting Beads, Invitrogen), and used to determine the number of viable cells per sample. Cells were stored at 4°C for 1-2 days and then run on a BD FACSymphony A5 High-Parameter Cell Analyzer. Beads (UltraComp eBeads, Invitrogen) stained with each individual antibody were used for compensation, with the exception of the viability dye control, where compensation was performed with leftover cells incubated with the viability dye alone. Data were analyzed in FlowJo_v10.8.1.

### Western Blotting

Mouse stomachs were aseptically harvested as described above and one third was frozen on dry ice and stored in liquid nitrogen. Tissues were lysed with a standard protocol with the indicated volumes of all reagents ^66^. Briefly, tissues were minced into fine pieces and then lysed in ice-cold RIPA buffer (Thermo Fisher) with added protease and phosphatase inhibitors (Thermo Fisher) using the gentleMACS Dissociator (Miltenyi). Protein concentrations were estimated with a Pierce BCA Protein Assay Kit (Thermo Fisher). Western blotting was performed as previously described ^67^. Densitometry was performed in ImageJ by inverting the pixel density for all blots, subtracting the background from each band to determine the net values, and calculating the ratio of net sample to net loading control.

### Organoids

Stomachs were aseptically harvested and opened along the lesser curvature, and food was gently removed using closed forceps. The forestomach and antrum were removed with a razor blade. Corpus tissue was then washed with 1xPBS. Tissues were cut into rice-sized pieces and cells were dissociated in 22.5 ml dissociation buffer (HBSS solution [Gibco] containing 0.01M HEPES, 1:100 Penicillin/Streptomycin, 1/100 L-glut, 0.00065M EDTA, 0.001M DTT) at 37°C for 35 minutes with gentle stirring. Cells were then filtered through a 100 μm strainer, washed with 1xPBS, and pelleted at 200xg for 5 minutes. The supernatant was removed and 10 ml neutralization buffer (HBSS with calcium and magnesium, Gibco) was added to quench the EDTA. Cells were again pelleted at 200xg for 5 minutes. The supernatant was removed and cells were resuspended in 200 μL ice cold Matrigel Basement Membrane Matrix (Corning) and plated into one well of a 6 well plate. Matrigel was allowed to solidify by incubating the plate at 37°C for 20 minutes, after which 3 mL of Mouse Intesticult OGM medium (StemCell) with 0.1 mg/ml Primocin (InvivoGen) was added to each well. Media was replaced every 2-3 days. After one to two passages, organoids were seeded to 40k cells per 60 μL of Matrigel into each well of a 12 well plate. Organoids were allowed to grow for 5 days, changing media every 2-3 days.

Three wells per organoid line were then fixed with 4% paraformaldehyde (PFA) for 30 minutes at room temperature while gently rocking. Organoids were suspended in 400 µl HistoGel (Fisher Scientific), solidified in disposable base molds (Sakura), and then processed for embedding and sectioning using standard histological protocols.

### Gastric cancer TMA

This study was approved by the Fred Hutchinson Cancer Center Institutional Review Board (IR8657). A database search was performed to identify subjects with gastric cancer who had tissues stored at the University of Washington Northwest BioSpecimen tissue repository, Seattle, WA, or stored at the Legacy Research Institute Tumor Bank, Portland, OR (the latter accessed through the Fred Hutch Specimen Acquisition Network, SAN) since 2010. Subjects were excluded if they had neoadjuvant therapy or insufficient stored materials. A pathologist (LK) reviewed the medical records to determine which tissue blocks to request. Available blocks were pulled and sectioned as follows: one 4 µm cut that was stained with hematoxylin and eosin (H&E), followed by three 10 µm unstained cuts, then one 4 µm H&E-stained cut. DNA was extracted from 1-2 unstained cuts using the AllPrep DNA/RNA FFPE kit (Qiagen). DNA concentrations were determined by spectrophotometry with a NanoDrop One. Twenty µl of DNA was used to perform droplet digital PCR to detect the *H. pylori* 16S rRNA gene as previously described ^68, 69^. Samples were tested in duplicate. Samples with 0 copies/µl DNA were considered negative. Samples with 1-2 copies/µl DNA were re-extracted and re-tested. The second H&E-stained cut was then annotated by a pathologist (CY) to note regions of superficial cancer, deep cancer and non-neoplastic tissue adjacent to cancer. The annotations were used to core the paraffin blocks and generate a tissue microarray (TMA) using TMA Grand Master (3DHISTECH) according to the manufacturer’s instructions. Thin sections (4 µm) were cut onto ProbeOn Plus slides (Fisher) and stained for immunofluorescence microscopy as described above.

### In situ hybridization and immunofluorescence microscopy

Immunofluorescence microscopy to detect gastric mucins, Ki-67, and tdTomato and GFP was performed on fixed tissue and organoid sections as previously described ^16^. Multiplex immune cell immunohistochemistry studies to detect macrophages and T cells were performed as previously described ^16^. *In situ* hybridization was performed on fixed tissue sections using the RNAscope system (ACD-biotechne) in accordance with the manufacturer’s instructions, using a Leica Bond RX autostainer. The RNAscope 2.5 LS Reagent Kit – BROWN was used for *Muc4* alone and the RNAscope LS Multiplex Fluorescent Assay Kit was used to assess multiple targets. Briefly, paraffin-embedded tissue sections were baked at 65°C for 60 minutes, then deparaffinized and hydrated on the Leica Bond RX autostainer. Heat-Induced Epitope Retrieval was performed with reagents from the indicated RNAscope reagent kits according to the manufacturer’s instructions. Briefly, tissues were incubated in Tris/EDTA, pH 9.0, for 15 minutes at 95°C, followed by a protein digestion in protease III for 15 minutes at room temperature. ISH probes and amplification reagents were applied according to the manufacturer’s instructions.

After staining, slides were manually dehydrated through graded alcohols, cleared in xylene, and mounted in Epredia Cytoseal XYL (Fisher). A list of antibodies, lectins and RNA probes is given in **Table S8**.

### Image Analysis

#### Analysis of mouse tissues

For multiplex immune cell immunohistochemistry studies, CD4, CD8α, and F4/80 were quantified as previously described using HALO software (Indica Labs) ^16^. *Muc4* ISH was scored in a blinded fashion using a semi-quantitative scale with the following criteria: 0 = no staining, 1 = 1-25% of corpus glands are positive, 2 = 26-50% of corpus glands are positive, 3 = 51-75% of corpus glands are positive, 4 = >75% of corpus glands are positive. For samples at the six week time point, the median score of 4-5 fields of view is reported due to greater heterogeneity in *Muc4* expression within individual mice. For lineage tracing studies of mouse stomach, 4-5 images of the corpus were taken and the total number of glands as well as the number of GFP+ glands were determined. For lineage tracing organoids, samples were prepared as described below and every organoid on the slide was inspected for GFP and tdTomato expression.

#### Analysis of human tissues

Immunostained TMA slides were scanned on a ScanScope FL slide scanner (Aperio Technologies) using the ScanScope Console.Ink software v120.0.0.33. Slide scans were converted to TIFF files for downstream analysis using Aperio Image Scope software v12.3.1.5011. The relationship between MUC4 and Ki-67 or MUC2 and MUC5Ac biomarkers in each TMA tissue core was quantified using a single cell resolution measurement providing fractional values of single or double positive cells from the total number of cells in each sample. TIFFs were imported into the open-source software QuPath 0.3.2 ^33^, and single cells were segmented from the DAPI (nuclear) signal using a built-in segmentation algorithm. Cell boundaries were approximated by tessellation of the nuclear masks, and the mean of either nuclear pixel intensity (for Ki-67) or whole cell pixel intensity (for mucins) was extracted. The percentage of single- or double- positive cells per TMA was then assessed. Because fluorescence intensity varied from slide to slide, analysis of marker abundance was conducted per slide. For each slide, the three tissue cores with the lowest average marker value were identified and visually inspected to confirm that they had low to no marker expression. These cores were used to set a baseline for negative/background staining. The average marker value and standard deviation was calculated for the three cores, and a threshold of five deviations above this baseline level was used as a cutoff for positive cells among all tissue cores on the given slide.

## Acknowledgements

The authors wish to acknowledge Melanie Dillon, Ciara Pike, Stephanie Weaver, Cassie Sathers and Feinan Wu for technical assistance. We thank Joseph Frascella, PhD, Vice President of Research and Carmen Rusinaru, MD, PhD, Senior Scientist at the Legacy Tumor Bank for providing gastric cancer tissue samples. This work was funded by an Innovation Grant from the Pathogen-Associated Malignancies Integrated Research Center (PAM-IRC) at Fred Hutchinson Cancer Center; a New Technology and Data Analysis Award from the PAM-IRC and Translational Data Science Integrated Research Center at Fred Hutchinson Cancer Center; a Gastric Cancer Foundation grant (DMG 2021-024 to NRS, MAK and SB); NIH R01 AI54423 and NIH R21 CA27051 (to NRS) and NIH K99 CA263036 (to VPO). Research was supported by the Cellular Imaging, Comparative Medicine, Genomics & Bioinformatics, Flow Cytometry and Research Pathology Shared Resources of the Fred Hutch/University of Washington Cancer Consortium (P30 CA015704). VPO was previously supported by an Irvington Postdoctoral Fellowship from the Cancer Research Institute, and was also supported by a Debbie’s Dream Foundation—AACR Gastric Cancer Research Fellowship, in memory of Sally Mandel (18-40- 41-OBRI).

## List of Supplemental Tables

Table S1. Description of the mouse samples used for single-cell RNA-sequencing.

Table S2. Genes driving the clustering of cells in UMAP #1; genes with power ≥ 0.4 are given.

Table S3. Genes driving the clustering of cells in UMAP #2. Up to the top 70 genes with the highest fold change and Padjusted < 0.05 are given for each cluster.

Table S4. The 10 most differentially expressed genes within each cluster of UMAP #2.

Table S5. Pathway analysis of metaplastic pit cells (pit_2, pit_6 and pit_8) versus classical pit cells (pit_1, pit_3, pit_4, pit_5 and pit_7) from UMAP #2.

Table S6. Characteristics of human subjects used in this study.

Table S7. Expression of mucins (MUC5Ac, MUC2 and MUC4) and the cell proliferation marker Ki-67 in each specimen in the gastric cancer tissue microarray (TMA). Table S8. List of antibodies, lectins and probes used in this study.

**Figure S1.**
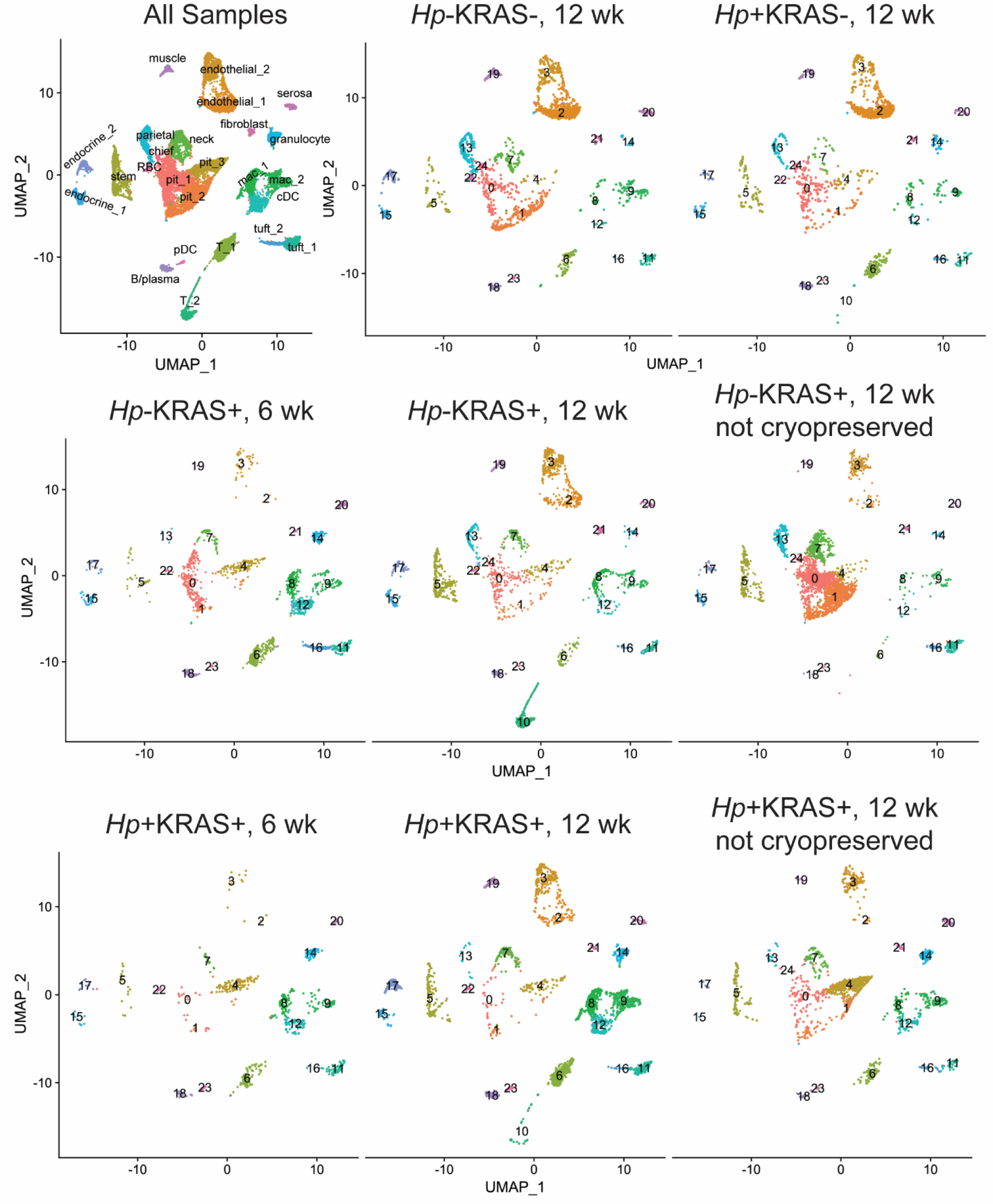
Summary of cell clusters detected in different mouse treatment groups at six and 12 weeks. Gastric single-cell RNA-sequencing was performed on eight samples obtained six or 12 weeks after *Hp* infection and/or constitutively active KRAS induction, and the UMAP from each sample is shown. Gastric single cell suspensions were prepared from mouse stomachs via digestion with a cold-active protease. Six of the samples were cryopreserved, then thawed, pooled as indicated in Table S1, subjected to dead cell removal and captured in Gel Beads in Emulsion (GEMs) for sequencing. Two samples, indicated as “not cryopreserved,” were captured in GEMs immediately following single cell suspension generation and dead cell removal. The two libraries from the six week timepoint were barcoded and sequenced in one run and the six libraries from the 12 week timepoint were barcoded and sequenced in a second run, as described in Table S1. After standard quality control metrics, the number of cells per sample ranged from 723 (*Hp*+KRAS+, 6 wk) to 3574 (*Hp*-KRAS+, 12 wk, not cryopreserved). The UMAP #1 from Figure 1B is reproduced in the top left, and the other UMAPs correspond to the indicated treatment groups. Clusters were annotated as follows: 0, pit cell_1; 1, pit cell_2; 2, endothelial cell_1; 3, endothelial cell_2; 4, pit cell_3; 5, stem cell; 6, T cell_1; 7, mucous neck cell; 8, macrophage_1; 9, macrophage_2; 10, T cell_2; 11, tuft cell_1; 12, conventional dendritic cell; 13, parietal cell; 14, granulocyte; 15, enteroendocrine cell_1; 16, tuft cell_2; 17, enteroendocrine cell_2; 18, B or plasma cell; 19, muscle cell; 20, serosal cell; 21, fibroblast; 22, erythrocyte or reticulocyte; 23, plasmacytoid dendritic cell; 24, chief cell.

**Figure S2.**
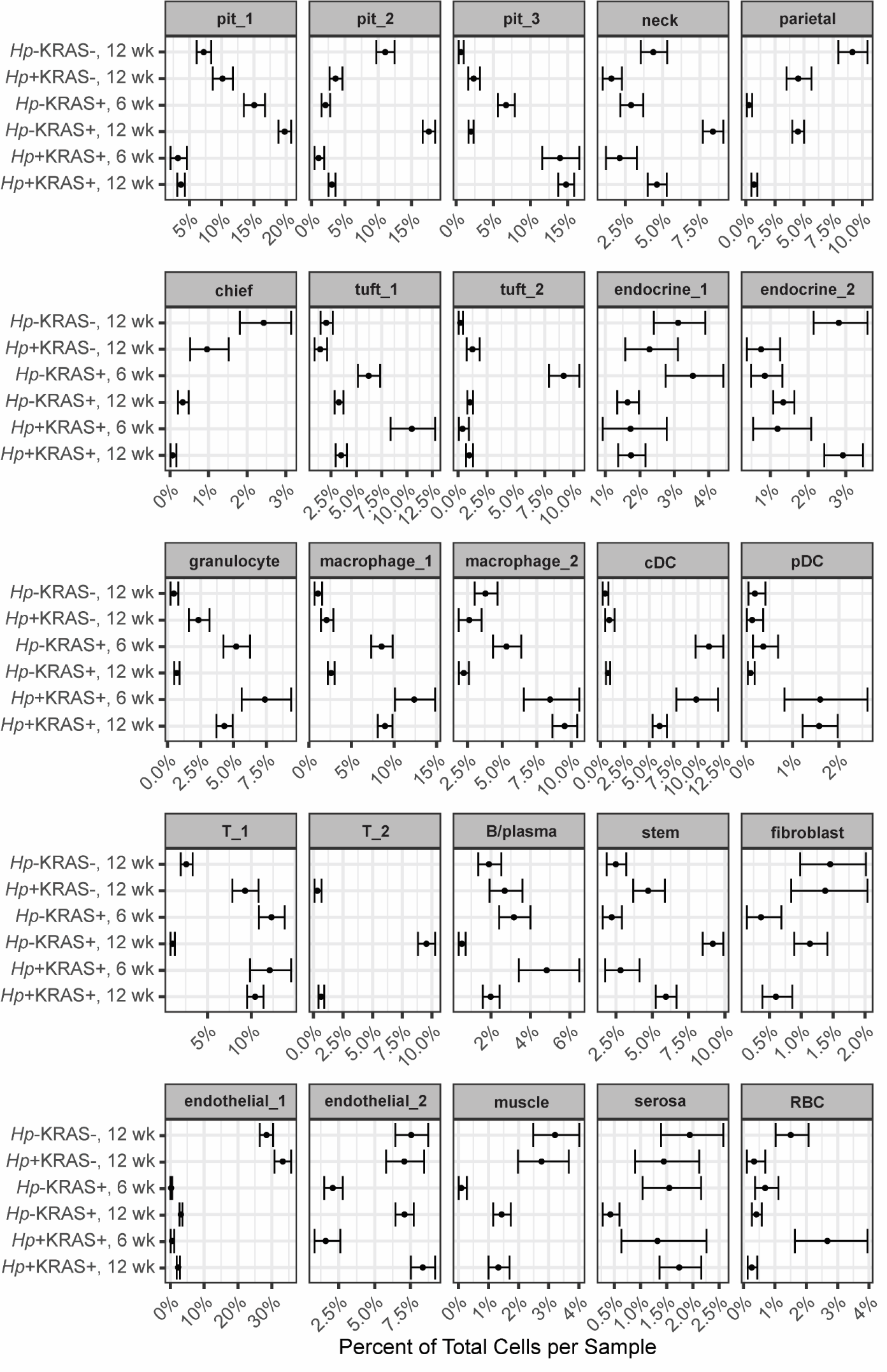
Cell cluster frequencies change according to *Hp* infection status and induction of constitutively active KRAS. The proportion of cells assigned to each annotated cluster in UMAP #1 is given. Samples were categorized according to treatment and time and the proportion and confidence level of each cell type was estimated from the empirical Bayesian distribution based on the observations of cell type occurrence using the EBBR package in R. Error bars represent the confidence interval that a given percentage of cells would be identified as the given type based on the distribution we observed. Endocrine, enteroendocrine cell; cDC, conventional dendritic cell; pDC, plasmacytoid dendritic cell, RBC, erythrocyte/reticulocyte.

**Figure S3.**
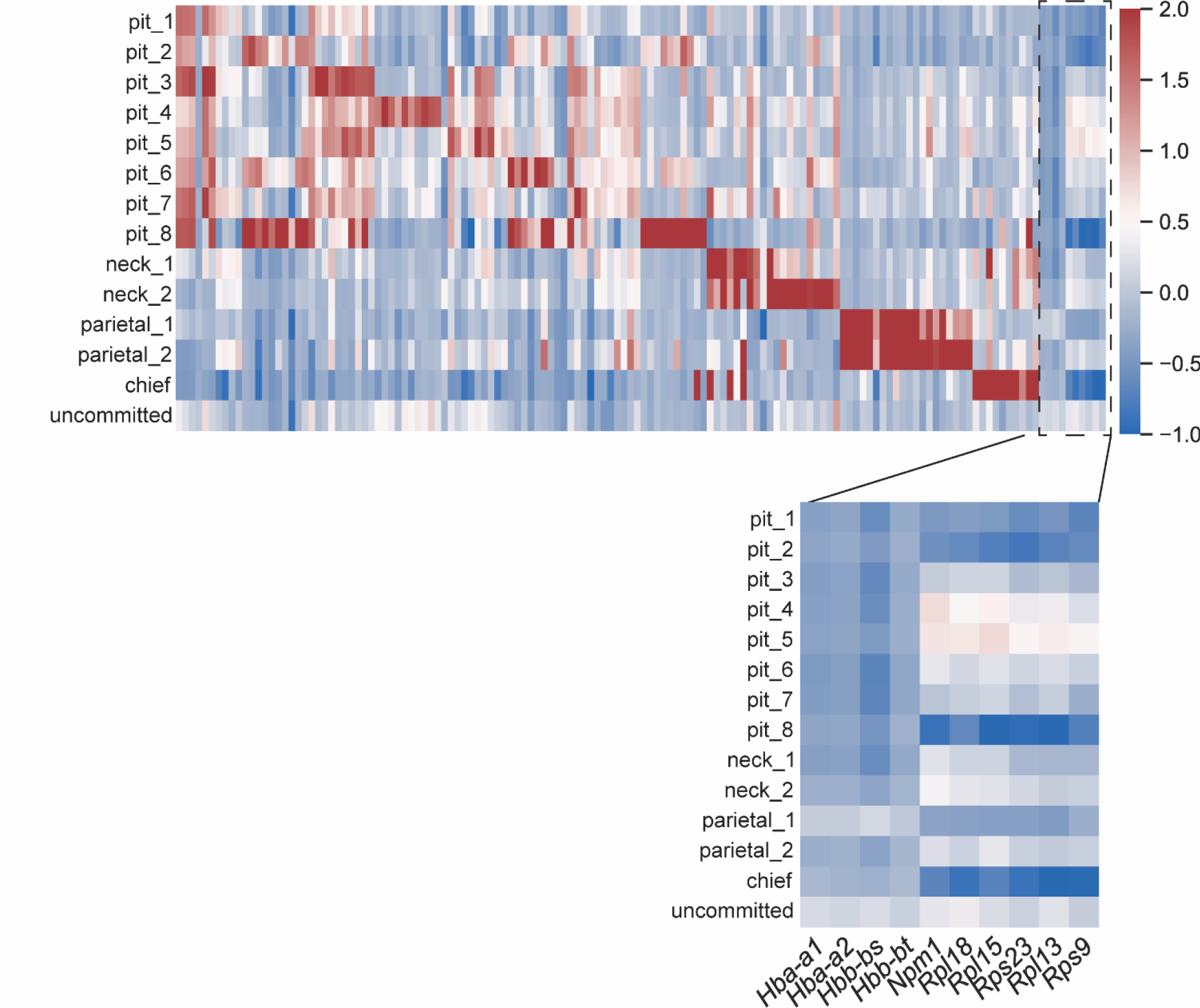
**Genes driving the subclustering of the major gastric epithelial cell types**. The heatmap shows the normalized expression of the top 10 genes driving the clustering of the indicated cell types from UMAP #2. The genes that segregate the “uncommitted” cluster (dashed line) are given. All genes in the heatmap are given in Table S4.

**Figure S4.**
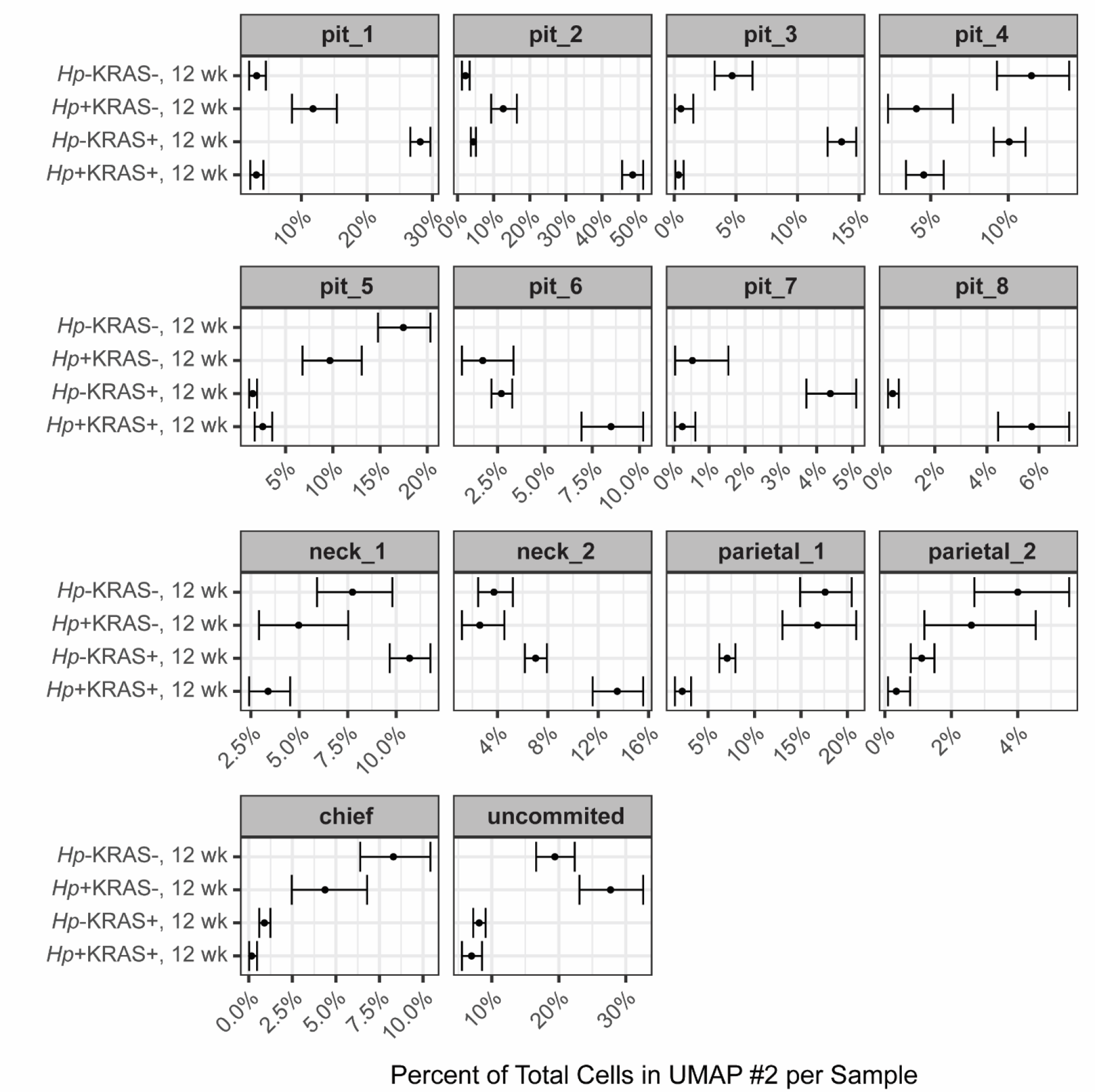
The epithelial subclusters pit_2, pit_6, pit_8 and neck_2 are expanded in *Hp*+KRAS+ mice. The proportion of cells assigned to each annotated cluster in UMAP #2 is given. Samples were categorized according to treatment and the proportion and confidence level of each cell type was estimated from the empirical Bayesian distribution based on the observations of cell type occurrence in R using the EBBR package. Error bars represent the confidence interval that a given percentage of cells would be identified as the given type based on the distribution we observed.

**Figure S5.**
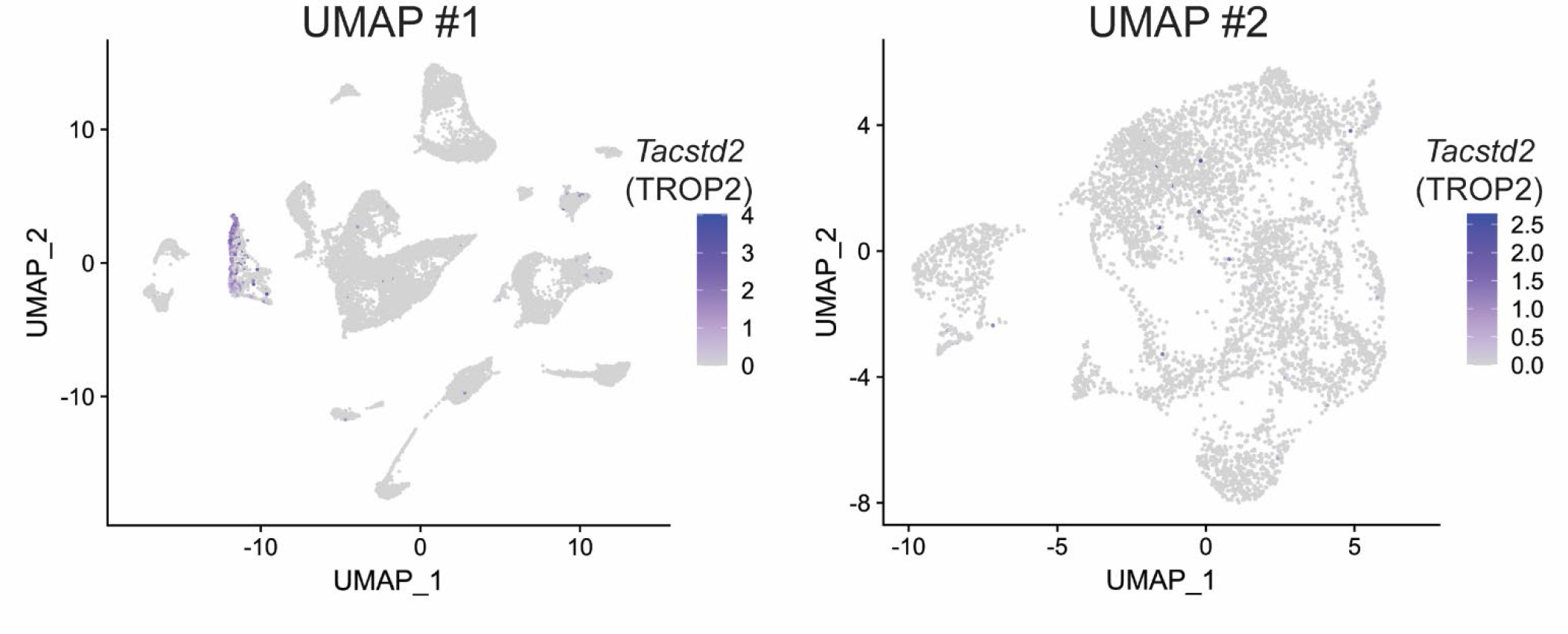
The dysplasia marker gene *Trop2* is rarely detected in the central epithelial megacluster. Cells expressing the indicated genes are highlighted in purple on UMAP #1 (left) and UMAP #2 (right). The color scale indicates the magnitude of gene expression within a cell, expressed as ln([count per 10,000 reads]+1).

**Figure S6.**
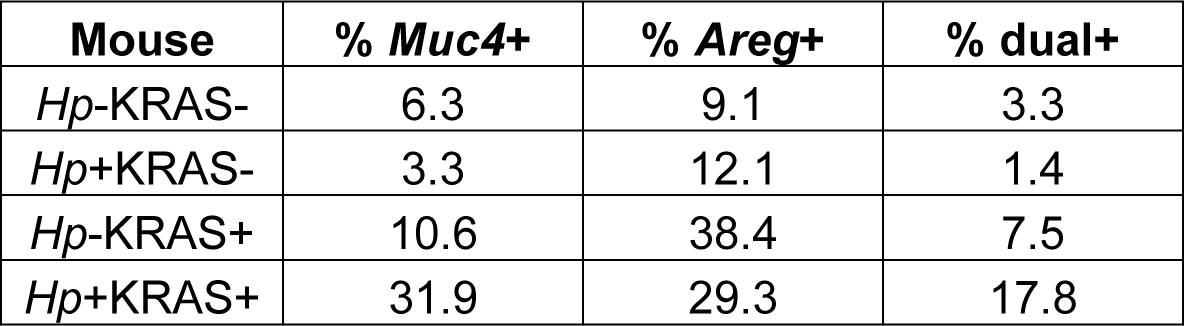
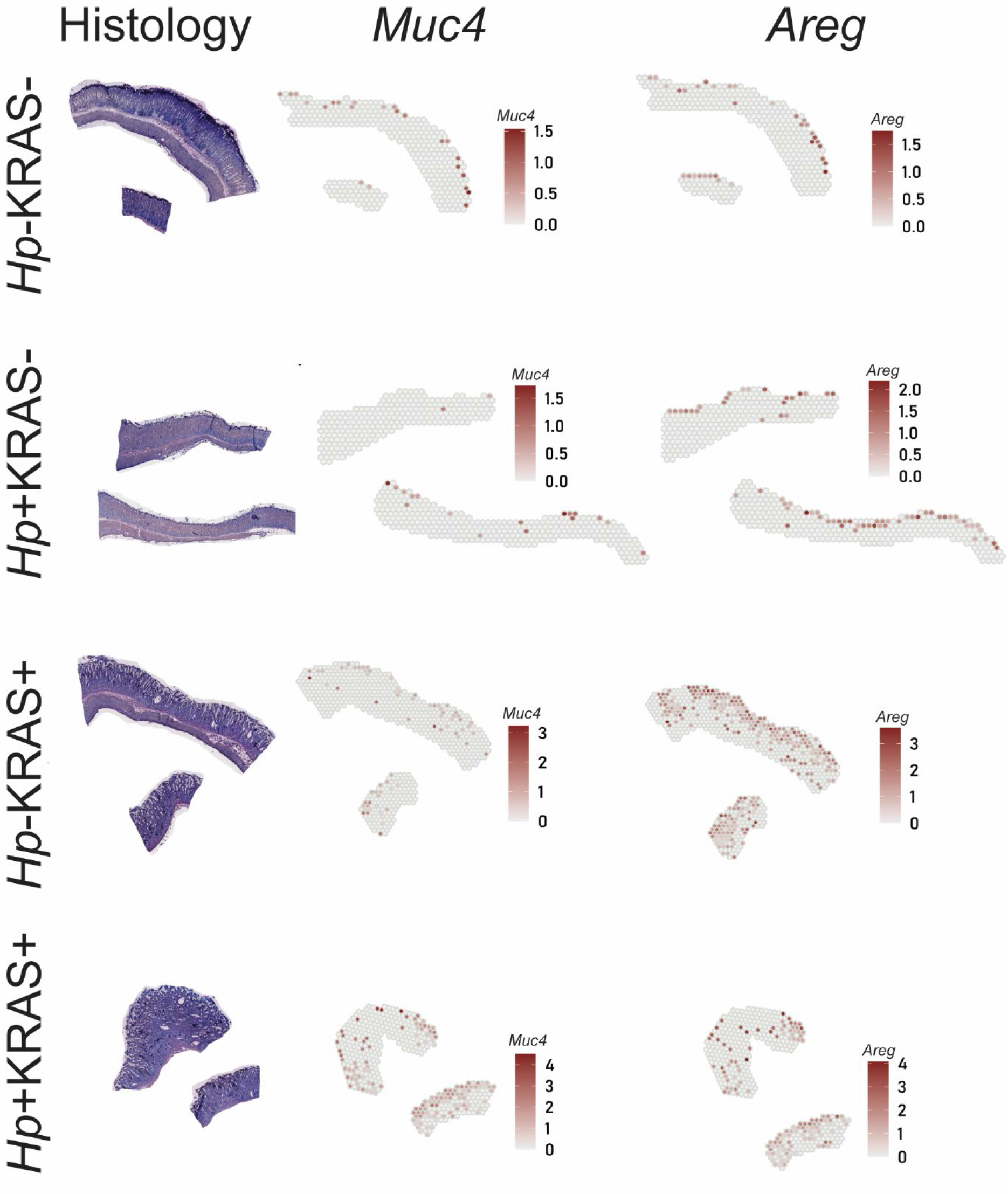
Spatial profiling reveals enrichment of *Muc4* and *Areg* expression from the luminal surface in the *Hp*+KRAS+ gastric epithelium. A spatial gene expression experiment conducted with the 10x Visium platform demonstrated greater metaplastic pit cell abundance in *Hp*+KRAS+ mice than in the other treatment groups. As described in the Supplemental Methods, at the 12 week time point, mice were humanely euthanized and tissues were cryo- preserved in OCT medium by plunge freezing in an isopentane/liquid nitrogen bath. Tissue sections were cut onto a barcoded Visium slide. After imaging, tissues were permeabilized to release RNA onto the barcoded spots. Libraries were prepared and sequenced and reads were mapped back to the barcoded spots using SpaceRanger (10x). Damaged and folded regions of tissue were omitted from the analysis. Spots that were positive for *Muc4* and *Areg* are indicated. The proportion of *Muc4*+, *Areg*+, and *Muc4*+*Areg*+ spots was:

**Figure S7.**
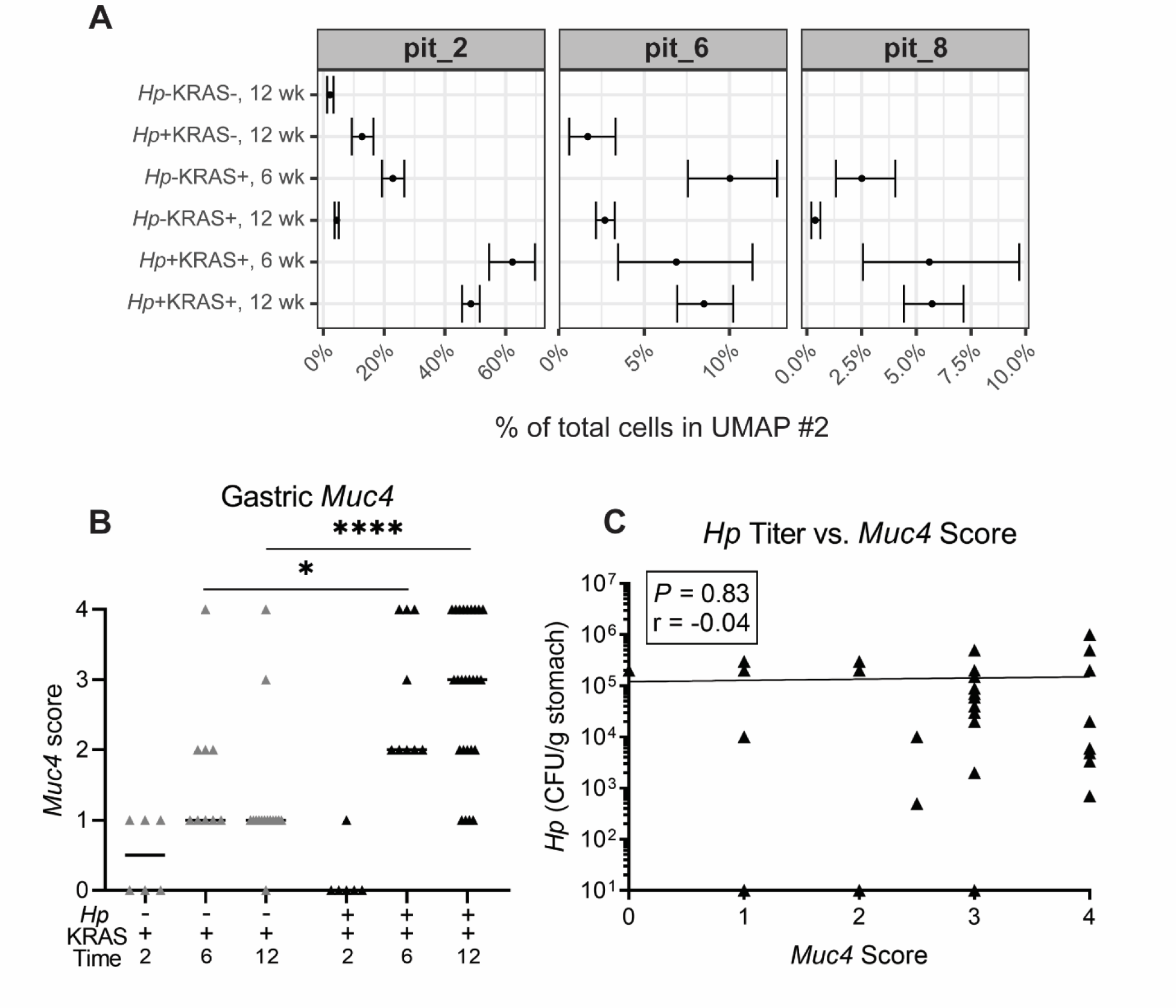
Metaplastic pit cells were found by six weeks and *Muc4* ISH score does not correlate with *Hp* titers. A) The proportion of cells assigned to the indicated pit cell subclusters is shown for both the six and 12 week time points. Error bars represent the confidence interval that a given percentage of cells would be identified as the given cluster type based on their observed distribution. **B)** Tissues were scored in a blinded fashion. N=5 independent experiments were performed with n=3-8 mice per group. Data represent actual values from each individual mouse and bars indicate the median values. **C)** The *Hp* stomach titer at six weeks is plotted against the *Muc4* ISH score from the same mouse. Statistical significance was assessed by a Spearman correlation and the trend line was fitted by a simple linear regression. CFU, colony-forming units.

**Figure S8.**
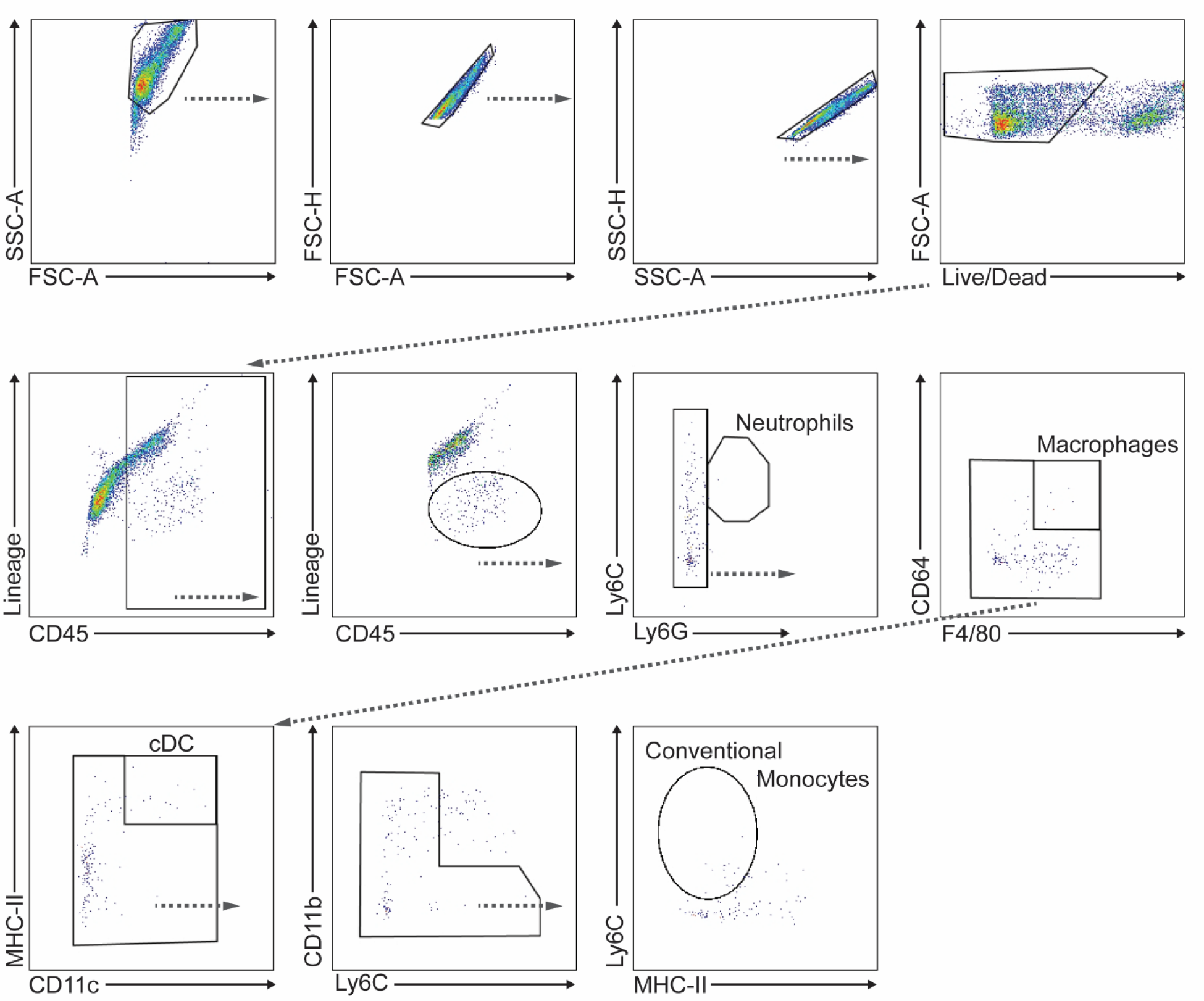
Gating strategy for detection of myeloid cell populations by flow cytometry. Mouse gastric lamina propria cells were isolated and stained with the indicated markers. A representative *Hp*+KRAS+ mouse is shown. ‘Lineage’ comprises B220, NK1.1 and CD90.2. cDC, conventional dendritic cell.

**Figure S9.**
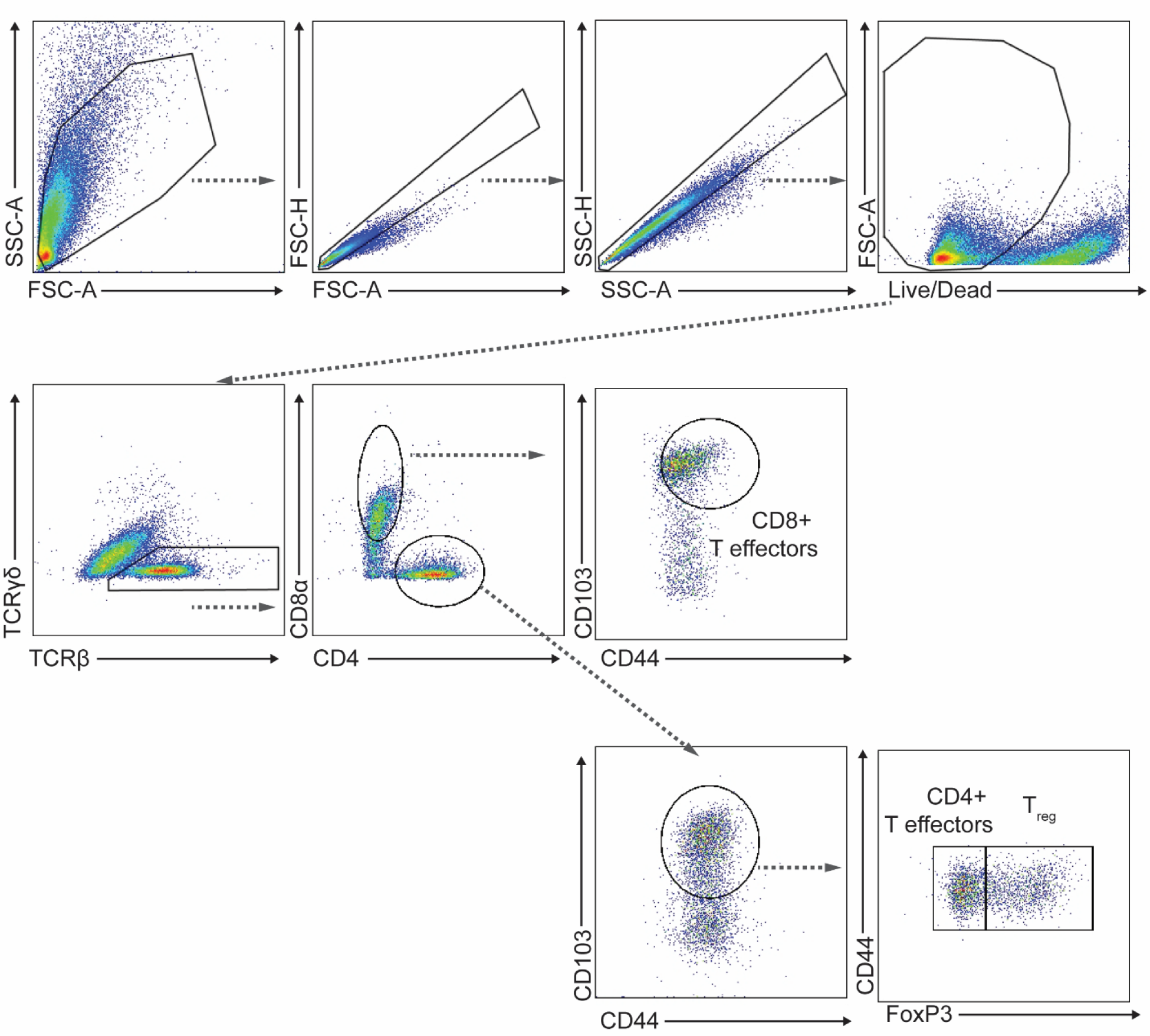
Gating strategy for detection of T cell populations by flow cytometry. Mouse gastric lamina propria cells were isolated and stained with the indicated markers. A representative *Hp*+KRAS+ mouse is shown.

**Figure S10.**
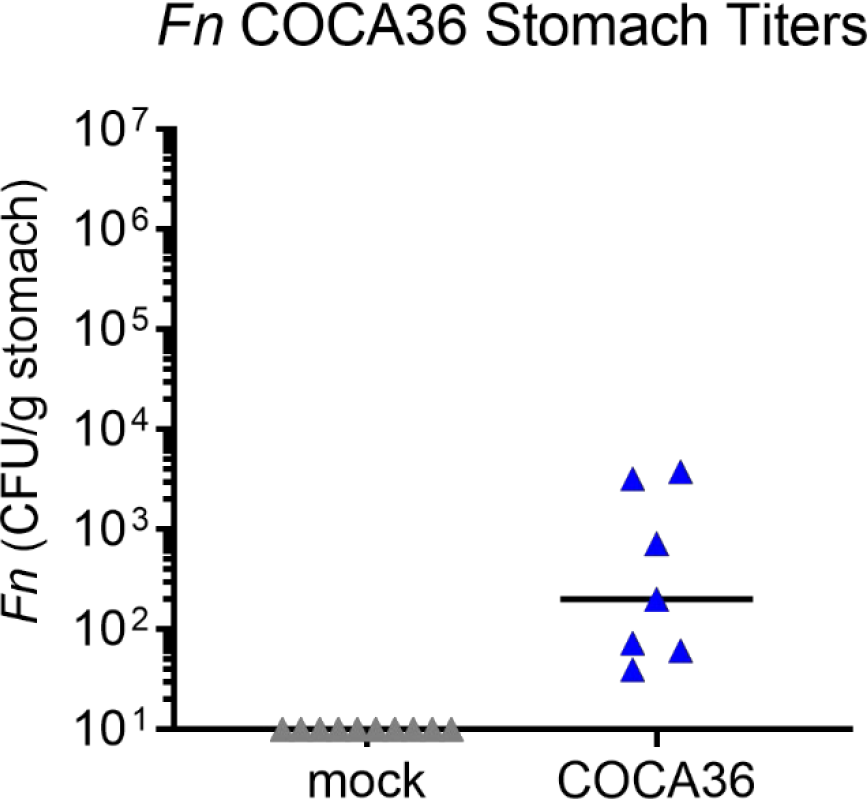
*Fusobacterium nucleatum* strain COCA36 is able to colonize the KRAS+ stomach. Mice were given tamoxifen to induce constitutively active KRAS expression. After six weeks, mice were mock-infected or infected with *F. nucleatum* strain COCA36. Shown are stomach titers at 12 weeks. Data are from N=2 independent experiments with n=3-5 mice per group. Data points represent actual values from individual mice and lines represent the median. Zeroes are plotted at the limit of detect (10 CFU).

**Figure S11.**
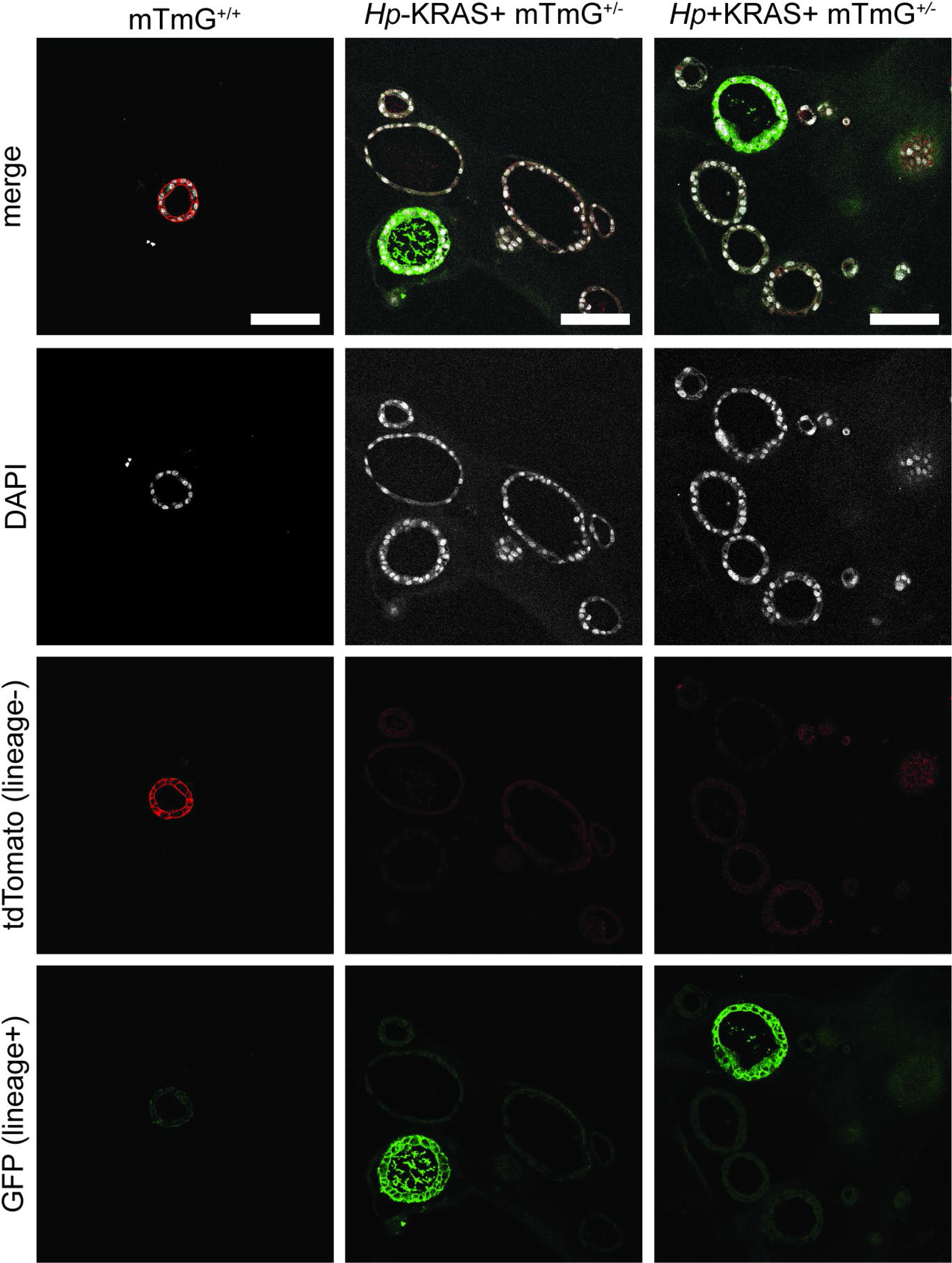
Few gastric organoids exhibit GFP staining indicative of lineage-derived cells. Organoids generated from lineage tracing mice were stained for tdTomato (lineage- negative glands), GFP (lineage-positive glands) and DAPI (nuclei). The same images from main body Figure 5 are shown and individual channels are given. Scale bars, 100 µm.

**Figure S12.**
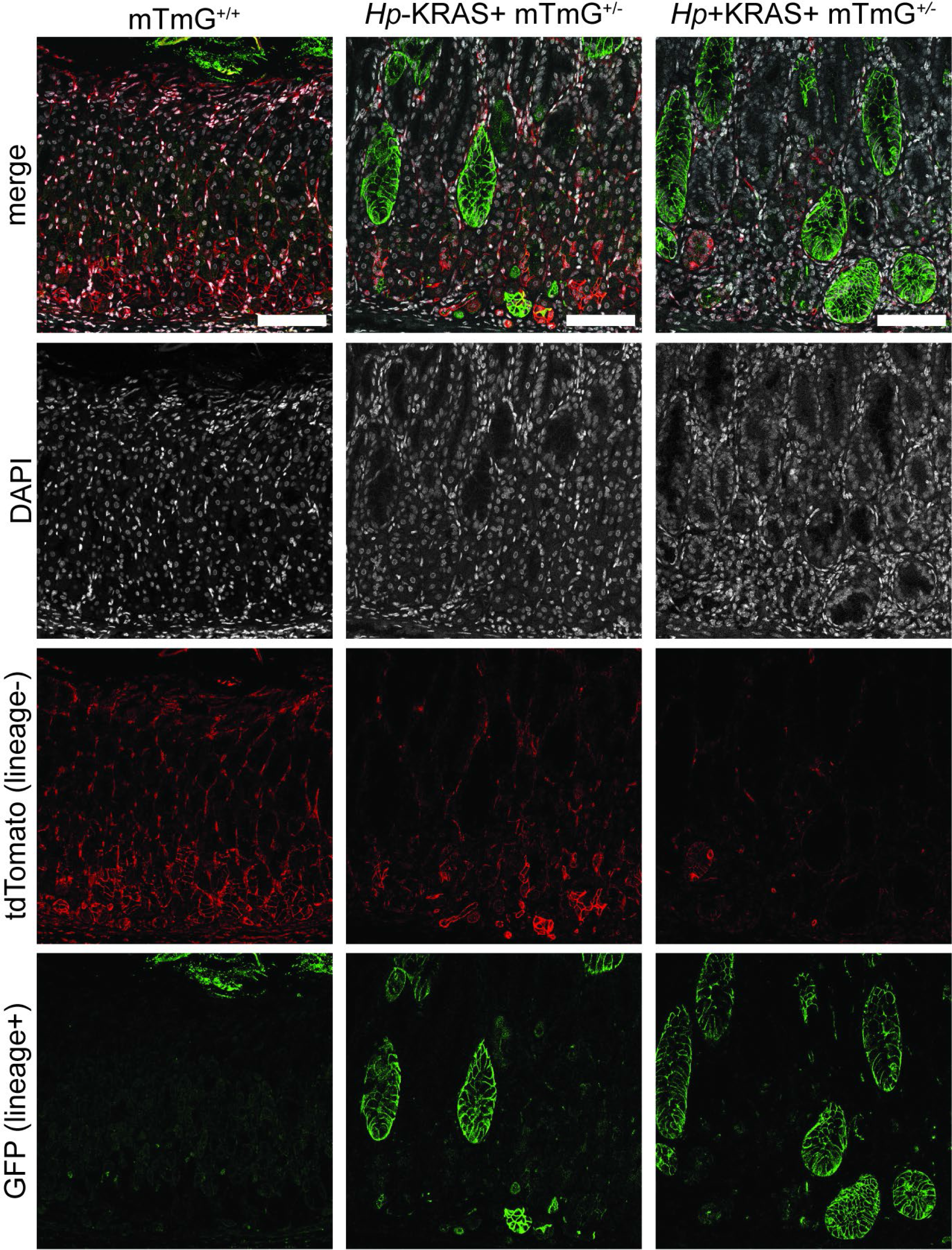
Few tdTomato+ glands are observed in lineage tracing mice, and *Hp*+KRAS+ mice have more GFP+ glands than *Hp*-KRAS+ mice do. Stomachs from lineage tracing mice were stained for tdTomato (lineage-negative glands), GFP (lineage-positive glands) and DAPI (nuclei). The same images from main body Figure 5 are shown and individual channels are given. Scale bars, 100 µm. The GFP signal at the top of the glands in the control mTmG^+/+^ mouse is autofluorescence from food contents in the stomach.

**Figure S13.**
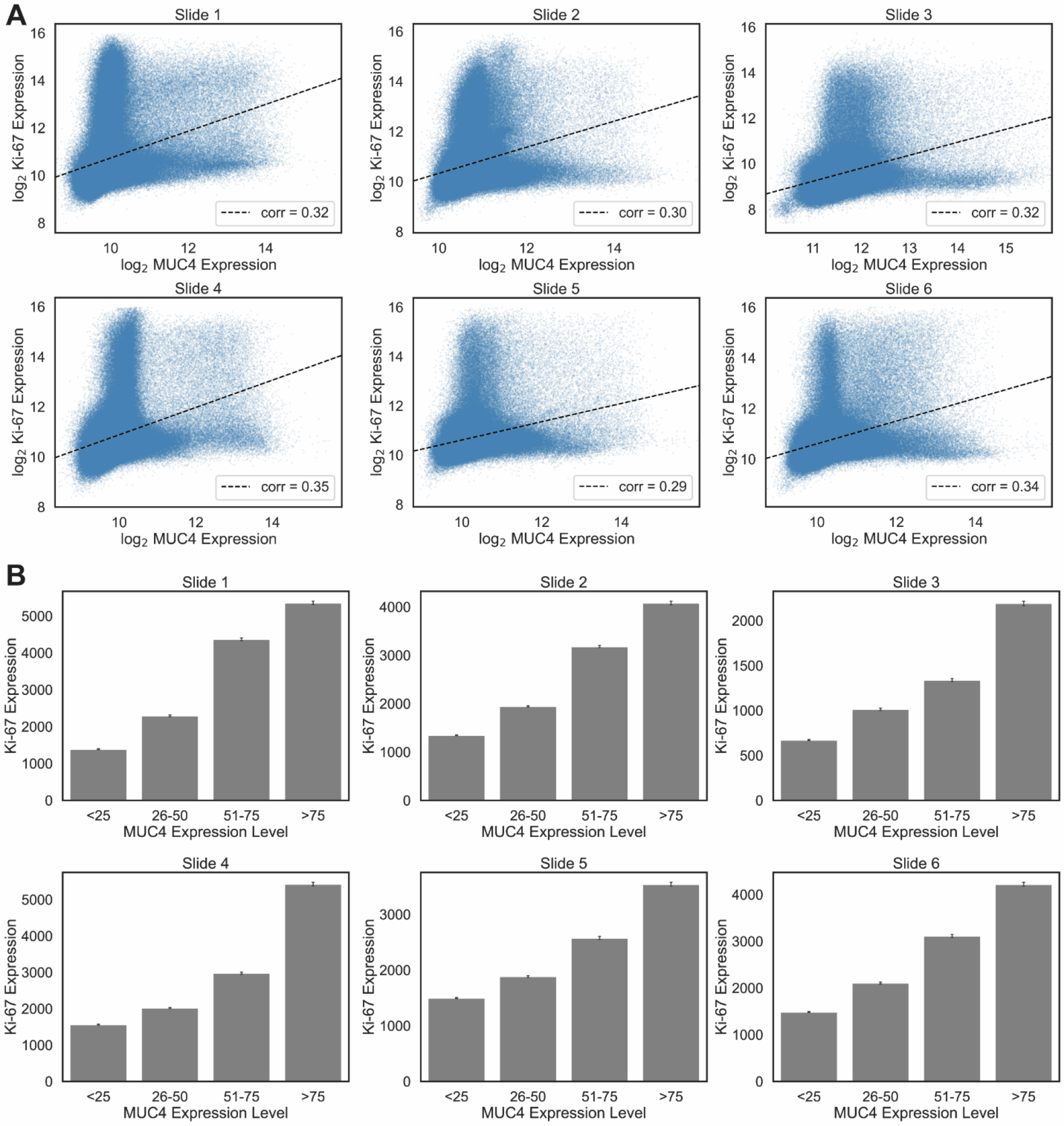
MUC4 and Ki-67 expression are significantly positively associated in samples from 47 subjects with gastric cancer. Samples were arranged in tumor microarrays (TMAs) comprising six paraffin blocks with 19-24 tissue cores per block. Each block was sectioned onto slides that were immunostained for MUC4 and Ki-67 expression and scanned for image analysis. Due to variability in staining intensity among the different slides, each slide was analyzed individually. QuPath was used to segment cells and detect marker expression within each cell of a given tissue core based on nuclear (Ki-67) and cellular (MUC4) signal intensity in the corresponding channels. **A)** The log_2_-transformed signal intensity of MUC4 and Ki-67 is plotted for every cell in every tissue core on each individual slide. The following numbers of cells were detected: slide 1, 506,369; slide 2, 477,241; slide 3, 344,500; slide 4, 499,623; slide 5, 438,598; slide 6, 424,600 (total n=2,690,991 cells). Statistical significance was tested with a Pearson’s correlation and the correlation coefficient (“corr”) is given; for each slide, *P* < 0.00001. For each slide, every cell in every tissue core was binned in quartiles according to its MUC4 expression level and the mean Ki-67 expression level of the cells in each quartile was calculated. Bars indicate the 95% confidence interval of the estimated true mean. Scatter and bar plots were made using the Seaborn package in Python.

**Figure S14.**
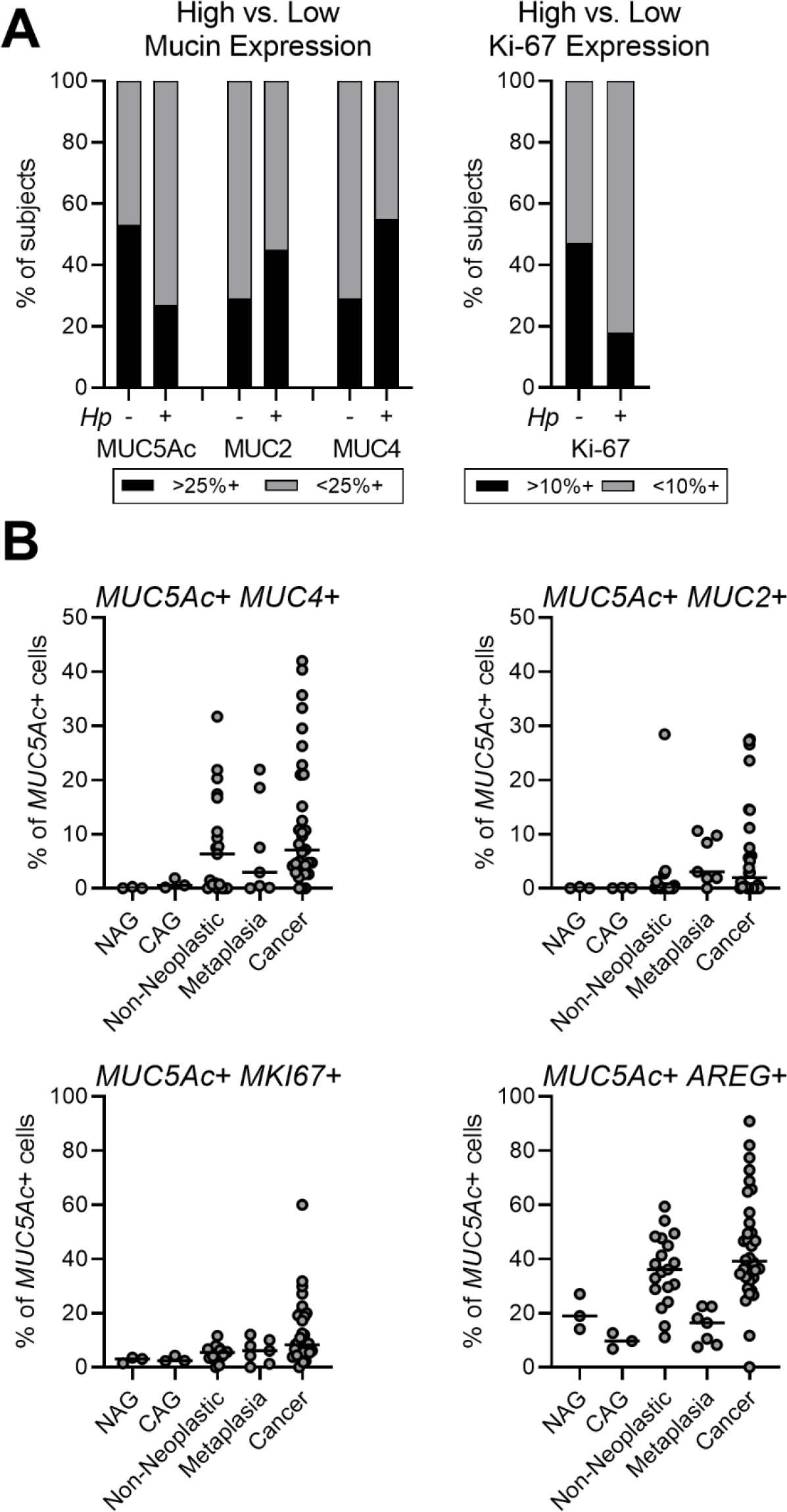
Pit cells can express intestinal mucins during preneoplasia and cancer. A) The data from **Figure 6E** are shown. The expression of the mucins MUC5Ac, MUC2 and MUC4 and the cell proliferation marker Ki-67 was probed in a tissue microarray (TMA) comprising samples from 47 gastric cancer (“CA”) patients. Gastric *Hp* status was determined by droplet digital PCR and subjects were categorized as *Hp*- or *Hp*+ to indicate whether their stomachs still harbored active *Hp* infection. For each subject, the tissue core with the greatest expression of each marker was assessed and the data are stratified according to whether expression was high (>25% mucin positivity or >10% Ki-67 positivity). **B)** Three published gastric scRNA-seq datasets were analyzed and the proportion of pit cells (*MUC5Ac*+) expressing the indicated disease markers is given. Each dot represents one scRNA-seq sample, which are arranged in order of disease severity. NAG, non-atrophic gastritis; CAG, chronic atrophic gastritis.

**Figure S15.**
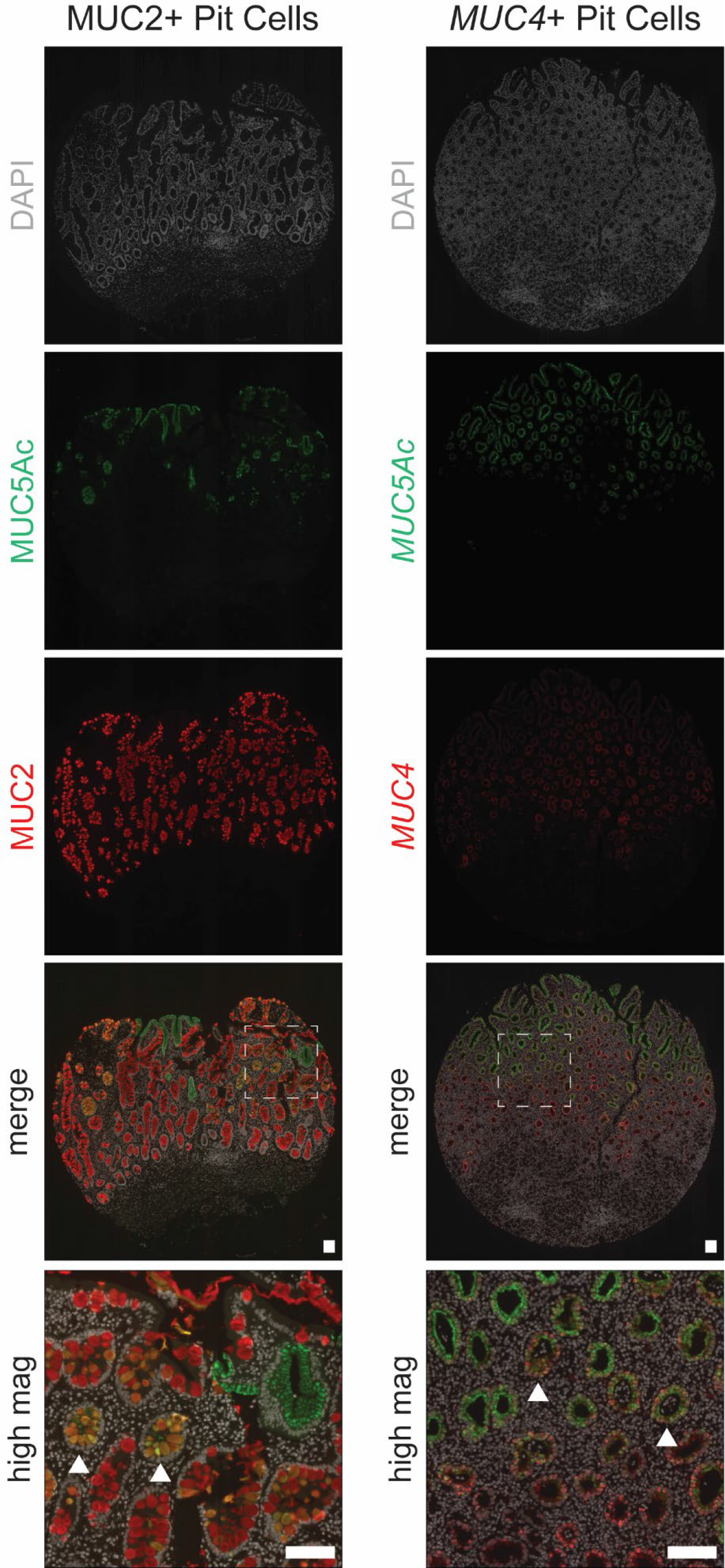
Pit cells can express the intestinal mucins MUC2 and MUC4. Shown are non- neoplastic epithelial samples from two different gastric cancer subjects. The left panel shows immunostaining for the pit cell mucin MUC5Ac (green) and the intestinal mucin MUC2 (red). The right panel shows *in situ* hybridization for *MUC5Ac* (green) and the intestinal mucin *MUC4* (red). The dashed boxes indicate regions shown at higher magnification below. Arrows show examples of glands containing dual-positive cells. Scale bars, 100 µm.

## References

1. Tlsty, T. D. & Gascard, P. Stromal directives can control cancer. Science 365, 122–123, doi:10.1126/science.aaw2368 (2019).

2. de Martel, C., Georges, D., Bray, F., Ferlay, J. & Clifford, G. M. Global burden of cancer attributable to infections in 2018: a worldwide incidence analysis. Lancet Glob Health, doi:10.1016/S2214-109X(19)30488-7 (2019).

3. Suerbaum, S. & Michetti, P. Helicobacter pylori infection. The New England journal of medicine 347, 1175–1186, doi:10.1056/NEJMra020542 (2002).

4. Choi, E., Hendley, A. M., Bailey, J. M., Leach, S. D. & Goldenring, J. R. Expression of Activated Ras in Gastric Chief Cells of Mice Leads to the Full Spectrum of Metaplastic Lineage Transitions. Gastroenterology 150, 918–930 e913, doi:10.1053/j.gastro.2015.11.049 (2016).

5. Caldwell, B., Meyer, A. R., Weis, J. A., Engevik, A. C. & Choi, E. Chief cell plasticity is the origin of metaplasia following acute injury in the stomach mucosa. Gut 71, 1068–1077, doi:10.1136/gutjnl-2021-325310 (2022).

6. Nam, K. T. et al. Mature chief cells are cryptic progenitors for metaplasia in the stomach. Gastroenterology 139, 2028–2037 e2029, doi:10.1053/j.gastro.2010.09.005 (2010).

7. Correa, P. A human model of gastric carcinogenesis. Cancer research 48, 3554–3560 (1988).

8. Goldenring, J. R., Nam, K. T., Wang, T. C., Mills, J. C. & Wright, N. A. Spasmolytic polypeptide-expressing metaplasia and intestinal metaplasia: time for reevaluation of metaplasias and the origins of gastric cancer. Gastroenterology 138, 2207–2210, 2210 e2201, doi:10.1053/j.gastro.2010.04.023 (2010).

9. Kusters, J. G., van Vliet, A. H. & Kuipers, E. J. Pathogenesis of Helicobacter pylori infection. Clinical microbiology reviews 19, 449–490, doi:10.1128/CMR.00054-05 (2006).

10. Lennerz, J. K. et al. The transcription factor MIST1 is a novel human gastric chief cell marker whose expression is lost in metaplasia, dysplasia, and carcinoma. The American journal of pathology 177, 1514–1533, doi:10.2353/ajpath.2010.100328 (2010).

11. Correa, P., Haenszel, W., Cuello, C., Tannenbaum, S. & Archer, M. A model for gastric cancer epidemiology. Lancet 2, 58–60 (1975).

12. Petersen, C. P. et al. Macrophages promote progression of spasmolytic polypeptide-expressing metaplasia after acute loss of parietal cells. Gastroenterology 146, 1727–1738 e1728, doi:10.1053/j.gastro.2014.02.007 (2014).

13. Kumar, S., Metz, D. C., Ellenberg, S., Kaplan, D. E. & Goldberg, D. S. Risk Factors and Incidence of Gastric Cancer After Detection of Helicobacter pylori Infection: A Large Cohort Study. Gastroenterology 158, 527–536 e527, doi:10.1053/j.gastro.2019.10.019 (2020).

14. Ford, A. C., Yuan, Y. & Moayyedi, P. Helicobacter pylori eradication therapy to prevent gastric cancer: systematic review and meta-analysis. Gut 69, 2113–2121, doi:10.1136/gutjnl-2020-320839 (2020).

15. Bang, C. S. et al. Helicobacter pylori Eradication for Prevention of Metachronous Recurrence after Endoscopic Resection of Early Gastric Cancer. J Korean Med Sci 30, 749–756, doi:10.3346/jkms.2015.30.6.749 (2015).

16. O’Brien, V. P. et al. Sustained Helicobacter pylori infection accelerates gastric dysplasia in a mouse model. Life Sci Alliance 4, doi:10.26508/lsa.202000967 (2021).

17. Quante, M., Marrache, F., Goldenring, J. R. & Wang, T. C. TFF2 mRNA transcript expression marks a gland progenitor cell of the gastric oxyntic mucosa. Gastroenterology 139, 2018–2027 e2012, doi:10.1053/j.gastro.2010.08.003 (2010).

18. Bockerstett, K. A. et al. Single-cell transcriptional analyses of spasmolytic polypeptide-expressing metaplasia arising from acute drug injury and chronic inflammation in the stomach. Gut 69, 1027–1038, doi:10.1136/gutjnl-2019-318930 (2020).

19. Bockerstett, K. A. et al. Single-Cell Transcriptional Analyses Identify Lineage-Specific Epithelial Responses to Inflammation and Metaplastic Development in the Gastric Corpus. Gastroenterology 159, 2116–2129 e2114, doi:10.1053/j.gastro.2020.08.027 (2020).

20. Lee, S. H. et al. Up-regulation of Aquaporin 5 Defines Spasmolytic Polypeptide-Expressing Metaplasia and Progression to Incomplete Intestinal Metaplasia. Cell Mol Gastroenterol Hepatol 13, 199–217, doi:10.1016/j.jcmgh.2021.08.017 (2022).

21. Riera, K. M. et al. Trop2 is upregulated in the transition to dysplasia in the metaplastic gastric mucosa. J Pathol 251, 336–347, doi:10.1002/path.5469 (2020).

22. Jonckheere, N. et al. The human mucin MUC4 is transcriptionally regulated by caudal-related homeobox, hepatocyte nuclear factors, forkhead box A, and GATA endodermal transcription factors in epithelial cancer cells. J Biol Chem 282, 22638–22650, doi:10.1074/jbc.M700905200 (2007).

23. O’Neil, A., Petersen, C. P., Choi, E., Engevik, A. C. & Goldenring, J. R. Unique Cellular Lineage Composition of the First Gland of the Mouse Gastric Corpus. J Histochem Cytochem 65, 47–58, doi:10.1369/0022155416678182 (2017).

24. Meyer, A. R. et al. Group 2 Innate Lymphoid Cells Coordinate Damage Response in the Stomach. Gastroenterology 159, 2077–2091 e2078, doi:10.1053/j.gastro.2020.08.051 (2020).

25. Bullman, S. et al. Analysis of Fusobacterium persistence and antibiotic response in colorectal cancer. Science 358, 1443–1448, doi:10.1126/science.aal5240 (2017).

26. Yamamura, K. et al. Fusobacterium nucleatum in gastroenterological cancer: Evaluation of measurement methods using quantitative polymerase chain reaction and a literature review. Oncol Lett 14, 6373–6378, doi:10.3892/ol.2017.7001 (2017).

27. Zaiss, D. M. W., Gause, W. C., Osborne, L. C. & Artis, D. Emerging functions of amphiregulin in orchestrating immunity, inflammation, and tissue repair. Immunity 42, 216–226, doi:10.1016/j.immuni.2015.01.020 (2015).

28. Muzumdar, M. D., Tasic, B., Miyamichi, K., Li, L. & Luo, L. A global double-fluorescent Cre reporter mouse. Genesis 45, 593–605, doi:10.1002/dvg.20335 (2007).

29. He, S., Nakada, D. & Morrison, S. J. Mechanisms of stem cell self-renewal. Annu Rev Cell Dev Biol 25, 377–406, doi:10.1146/annurev.cellbio.042308.113248 (2009).

30. Radtke, F. & Clevers, H. Self-renewal and cancer of the gut: two sides of a coin. Science 307, 1904-1909, doi:10.1126/science.1104815 (2005).

31. Fiocca, R. et al. The foveolar cell component of gastric cancer. Hum Pathol 21, 260–270, doi:10.1016/0046-8177(90)90225-t (1990).

32. Rakic, S., Bandovic, J., Dunjic, M. & Randjelovic, T. Gastric foveolar hyperplasia in patients with cancer of the intact stomach. Surg Laparosc Endosc 4, 196–199 (1994).

33. Bankhead, P. et al. QuPath: Open source software for digital pathology image analysis. Sci Rep 7, 16878, doi:10.1038/s41598-017-17204-5 (2017).

34. Tang, Y. L., Gan, R. L., Dong, B. H., Jiang, R. C. & Tang, R. J. Detection and location of Helicobacter pylori in human gastric carcinomas. World J Gastroenterol 11, 1387–1391, doi:10.3748/wjg.v11.i9.1387 (2005).

35. Talarico, S. et al. Increased H. pylori stool shedding and EPIYA-D cagA alleles are associated with gastric cancer in an East Asian hospital. PLoS One 13, e0202925, doi:10.1371/journal.pone.0202925 (2018).

36. Cristescu, R. et al. Molecular analysis of gastric cancer identifies subtypes associated with distinct clinical outcomes. Nature medicine 21, 449–456, doi:10.1038/nm.3850 (2015).

37. Zhang, P. et al. Dissecting the Single-Cell Transcriptome Network Underlying Gastric Premalignant Lesions and Early Gastric Cancer. Cell Rep 27, 1934–1947 e1935, doi:10.1016/j.celrep.2019.04.052 (2019).

38. Sathe, A. et al. Single-Cell Genomic Characterization Reveals the Cellular Reprogramming of the Gastric Tumor Microenvironment. Clin Cancer Res 26, 2640–2653, doi:10.1158/1078-0432.CCR-19-3231 (2020).

39. Kumar, V. et al. Single-Cell Atlas of Lineage States, Tumor Microenvironment, and Subtype-Specific Expression Programs in Gastric Cancer. Cancer Discov 12, 670–691, doi:10.1158/2159-8290.CD-21-0683 (2022).

40. Zhang, C. T., He, K. C., Pan, F., Li, Y. & Wu, J. Prognostic value of Muc5AC in gastric cancer: A meta-analysis. World J Gastroenterol 21, 10453–10460, doi:10.3748/wjg.v21.i36.10453 (2015).

41. van den Brink, G. R. et al. Sonic hedgehog expression correlates with fundic gland differentiation in the adult gastrointestinal tract. Gut 51, 628–633, doi:10.1136/gut.51.5.628 (2002).

42. Lopez-Ferrer, A. et al. Role of fucosyltransferases in the association between apomucin and Lewis antigen expression in normal and malignant gastric epithelium. Gut 47, 349–356, doi:10.1136/gut.47.3.349 (2000).

43. Senapati, S. et al. Deregulation of MUC4 in gastric adenocarcinoma: potential pathobiological implication in poorly differentiated non-signet ring cell type gastric cancer. Br J Cancer 99, 949–956, doi:10.1038/sj.bjc.6604632 (2008).

44. Ganguly, K., Rauth, S., Marimuthu, S., Kumar, S. & Batra, S. K. Unraveling mucin domains in cancer and metastasis: when protectors become predators. Cancer Metastasis Rev 39, 647–659, doi:10.1007/s10555-020-09896-5 (2020).

45. Berasain, C. & Avila, M. A. Amphiregulin. Semin Cell Dev Biol 28, 31–41, doi:10.1016/j.semcdb.2014.01.005 (2014).

46. Nam, K. T. et al. Amphiregulin-deficient mice develop spasmolytic polypeptide expressing metaplasia and intestinal metaplasia. Gastroenterology 136, 1288–1296, doi:10.1053/j.gastro.2008.12.037 (2009).

47. Wang, B. et al. Abnormal amphiregulin expression correlates with gastric cancer prognosis. Oncotarget 7, 76684–76692, doi:10.18632/oncotarget.12436 (2016).

48. Jiang, J. et al. Over expression of amphiregulin promoted malignant progression in gastric cancer. Pathol Res Pract 215, 152576, doi:10.1016/j.prp.2019.152576 (2019).

49. Hayakawa, Y. et al. Mist1 Expressing Gastric Stem Cells Maintain the Normal and Neoplastic Gastric Epithelium and Are Supported by a Perivascular Stem Cell Niche. Cancer Cell 28, 800–814, doi:10.1016/j.ccell.2015.10.003 (2015).

50. Choi, E., Means, A. L., Coffey, R. J. & Goldenring, J. R. Active Kras Expression in Gastric Isthmal Progenitor Cells Induces Foveolar Hyperplasia but Not Metaplasia. Cell Mol Gastroenterol Hepatol 7, 251–253 e251, doi:10.1016/j.jcmgh.2018.09.007 (2019).

51. Matsuo, J. et al. Identification of Stem Cells in the Epithelium of the Stomach Corpus and Antrum of Mice. Gastroenterology 152, 218–231 e214, doi:10.1053/j.gastro.2016.09.018 (2017).

52. Kinoshita, H. et al. Three types of metaplasia model through Kras activation, Pten deletion, or Cdh1 deletion in the gastric epithelium. J Pathol 247, 35-47, doi:10.1002/path.5163 (2019).

53. Mueller, A., Merrell, D. S., Grimm, J. & Falkow, S. Profiling of microdissected gastric epithelial cells reveals a cell type-specific response to Helicobacter pylori infection. Gastroenterology 127, 1446–1462, doi:10.1053/j.gastro.2004.08.054 (2004).

54. Wroblewski, L. E. et al. Targeted mobilization of Lrig1(+) gastric epithelial stem cell populations by a carcinogenic Helicobacter pylori type IV secretion system. Proc Natl Acad Sci U S A 116, 19652–19658, doi:10.1073/pnas.1903798116 (2019).

55. Andrianifahanana, M. et al. IFN-gamma-induced expression of MUC4 in pancreatic cancer cells is mediated by STAT-1 upregulation: a novel mechanism for IFN-gamma response. Oncogene 26, 7251–7261, doi:10.1038/sj.onc.1210532 (2007).

56. Perrais, M. et al. Characterization of human mucin gene MUC4 promoter: importance of growth factors and proinflammatory cytokines for its regulation in pancreatic cancer cells. J Biol Chem 276, 30923–30933, doi:10.1074/jbc.M104204200 (2001).

57. Sayi, A. et al. The CD4+ T cell-mediated IFN-gamma response to Helicobacter infection is essential for clearance and determines gastric cancer risk. J Immunol 182, 7085–7101, doi:10.4049/jimmunol.0803293 (2009).

58. Farrow, D. C. et al. Use of aspirin and other nonsteroidal anti-inflammatory drugs and risk of esophageal and gastric cancer. Cancer epidemiology, biomarkers & prevention : a publication of the American Association for Cancer Research, cosponsored by the American Society of Preventive Oncology 7, 97–102 (1998).

59. Cancer Genome Atlas Research, N. Comprehensive molecular characterization of gastric adenocarcinoma. Nature 513, 202-209, doi:10.1038/nature13480 (2014).

60. Mejias-Luque, R. et al. Inflammation modulates the expression of the intestinal mucins MUC2 and MUC4 in gastric tumors. Oncogene 29, 1753–1762, doi:10.1038/onc.2009.467 (2010).

61. Arnold, I. C. et al. Tolerance rather than immunity protects from Helicobacter pylori- induced gastric preneoplasia. Gastroenterology 140, 199–209, doi:10.1053/j.gastro.2010.06.047 (2011).

62. Potter, A. S. & Steven Potter, S. Dissociation of Tissues for Single-Cell Analysis. Methods Mol Biol 1926, 55–62, doi:10.1007/978-1-4939-9021-4_5 (2019).

63. Subramanian, A. et al. Gene set enrichment analysis: a knowledge-based approach for interpreting genome-wide expression profiles. Proc Natl Acad Sci U S A 102, 15545–15550, doi:10.1073/pnas.0506580102 (2005).

64. Liberzon, A. et al. Molecular signatures database (MSigDB) 3.0. Bioinformatics 27, 1739–1740, doi:10.1093/bioinformatics/btr260 (2011).

65. Zhao, E. et al. Spatial transcriptomics at subspot resolution with BayesSpace. Nat Biotechnol 39, 1375–1384, doi:10.1038/s41587-021-00935-2 (2021).

66. Anderson K. B. B. Protein extraction and western blot (mouse tissues).

67. Sichel, S. R., Bratton, B. P. & Salama, N. R. Distinct regions of H. pylori’s bactofilin CcmA regulate protein-protein interactions to control helical cell shape. Elife 11, doi:10.7554/eLife.80111 (2022).

68. Talarico, S. et al. Quantitative Detection and Genotyping of Helicobacter pylori from Stool using Droplet Digital PCR Reveals Variation in Bacterial Loads that Correlates with cagA Virulence Gene Carriage. Helicobacter 21, 325–333, doi:10.1111/hel.12289 (2016).

69. Talarico, S. et al. High prevalence of Helicobacter pylori clarithromycin resistance mutations among Seattle patients measured by droplet digital PCR. Helicobacter 23, e12472, doi:10.1111/hel.12472 (2018).

