## Supplementary material for "Single-cell profiling uncovers a *Muc4*-expressing metaplastic gastric cell type sustained by *Helicobacter pylori*-specific inflammation": Table S8

**Table S8. List of antibodies, lectins and probes used in this study.**

Immunohistochemistry of mouse tissues and organoids:

| **Marker** | **Species** | **Dilution** | **Source** | **Purpose** |
| --- | --- | --- | --- | --- |
| Ki-67 | Rabbit | 1:300 | 12202; Cell Signaling Technology | Proliferating cell marker |
| RFP | Rabbit | 1:75 (stomachs), 1:150 (organoids) | 600-401-379S, Rockland | Shows lineage-negative (tdTomato+) glands |
| GFP | Chicken | 1:1000 | A10262, Fisher | Shows lineage-positive glands |

Immunohistochemistry of human tissues:

| **Marker** | **Species** | **Dilution** | **Source** | **Purpose** |
| --- | --- | --- | --- | --- |
| Ki-67 | Rabbit | 1:300 | 12202; Cell Signaling Technology | Proliferating cell marker |
| MUC4 | Mouse | 1:1000 | 8G7, Surinder K. Batra (UNMC) | Metaplasia marker |
| MUC2 | Rabbit | 1:300 | sc-15334; Santa Cruz | Metaplasia marker |
| MUC5Ac | Mouse | 1:500 | 45M1; Invitrogen | Gastric mucin |

Secondary antibodies used at 1:500 dilution:

| **Host** | **Reactive Against** | **Flour** | **Source** |
| --- | --- | --- | --- |
| Donkey | Rabbit | AF488, AF647 | Invitrogen |
| Donkey | Mouse | AF594, AF647 | Invitrogen |
| Goat | Chicken | AF488 | Invitrogen |

RNAscope® 2.5 LS probes from ACD-biotechne used for *in situ* hybridization:

| **Target** | **Catalog Number** |
| --- | --- |
| Mouse *Muc4* | 534018 |
| Mouse *Areg* | 430508-C3 |
| Human *MUC4* | 312888 |
| Human *MUC5Ac* | 312898-C2 |

Antibodies and Opal dyes used for mouse immune cell immunohistochemistry:

| **Position** | **Antibody** | **Clone & Host** | **Manufacturer & Catalog Number** | **Dilution**  **(Conc.)** | **Secondary** | **Opal Dye** |
| --- | --- | --- | --- | --- | --- | --- |
| 1 | CD3 | SP7 Rabbit | Thermo  RM-9107-S | 1:400 | Powervision Rabbit-HRP | **Opal 520** |
| 2 | F4/80 | D2S9R Rabbit | Cell Signaling  20076S | 1:4000  (0.1 µg/ml) | Powervision Rabbit-HRP | **Opal 540** |
| 3 | CD4 | 4SM95  Rat | eBioscience  14-9766-32 | 1:250  (2 µg/mL) | ImmPress Rat-HRP | **Opal 570** |
| 4 | CD8α | 4SM15  Rat | eBioscience  14-0808-82 | 1:1000  (0.5 µg/ml) | ImmPress Rat-HRP | **Opal 780** |

Antibodies used for Western blotting:

| **Target** | **Clone & Host** | **Manufacturer & Catalog Number** | **Dilution**  **in 10 ml** | **Secondary** |
| --- | --- | --- | --- | --- |
| EGFR | EP38Y rabbit | Abcam  ab52894 | 1:750 | goat anti-rabbit IgG-HRP (Santa Cruz) |
| phospho-EGFR (Y1068) | D7A5 rabbit | Cell Signaling  3777 | 1:1000 | goat anti-rabbit IgG-HRP  (Santa Cruz) |
| alpha-tubulin | DM1A  mouse | Cell Signaling  3873 | 1:2000 | goat anti-mouse IgG-HRP  (Santa Cruz) |

Antibodies used for flow cytometry:

| **Target** | **Fluor** | **Clone** | **Catalog Number** | **Manufacturer** | **Dilution** |
| --- | --- | --- | --- | --- | --- |
| B220 | BV510 | RA3-6B2 | 103248 | BioLegend | 1:200 |
| CD103 | A488 | 2E7 | 121420 | BioLegend | 1:300 |
| CD103 | BUV737 | M290 | 741739 | BD | 1:200 |
| CD11b | BV605 | M1/70 | 101257 | BioLegend | 1:400 |
| CD11c | PeCy7 | N418 | 25-0114-82 | eBioscience | 1:300 |
| CD4 | PC-eF710 | GK1.5 | 46-0041-82 | eBioscience | 1:200 |
| CD44 | AF700 | IM7 | 103026 | BioLegend | 1:200 |
| CD45 | BUV395 | 30-F11 | 564279 | BD | 1:250 |
| CD64 | PE | X54-5/7.1 | 139304 | BioLegend | 1:64 |
| CD8α | BUV395 | 53-6.7 | 563786 | BD | 1:200 |
| CD90.2 | BV510 | 30-H12 | 105335 | BioLegend | 1:200 |
| F4/80 | BV650 | T45-2342 | 743282 | BD | 1:200 |
| FoxP3 | BV421 | FJK-16s | 48-5773-82 | eBioscience | 1:150 |
| Ly6C | BV711 | HK1.4 | 128037 | BioLegend | 1:400 |
| Ly6G | BUV563 | 1A8 | 612921 | BD | 1:400 |
| MHC-II | PC-eF710 | M5/114.15.2 | 46-5321-82 | eBioscience | 1:500 |
| NK1.1 | BV510 | PK136 | 108738 | BioLegend | 1:400 |
| TCRβ | BV650 | H57-597 | 109251 | BioLegend | 1:200 |
| TCRγδ | FITC | GL3 | 118128 | BioLegend | 1:200 |
